# Frequency-selective oscillatory control of working memory robustness to distractors

**DOI:** 10.1101/2020.12.13.422600

**Authors:** Nikita Novikov, Boris Gutkin

## Abstract

Working memory (WM) is the brain’s ability to retain information that is not directly available from the sensory systems. WM retention is accompanied by sustained firing rate modulation and changes of the large-scale oscillatory profile. Among other changes, beta-band activity elevates in task-related regions, presumably stabilizing WM retention. Alpha-band activity, in turn, is stronger in task-irrelevant regions, serving to protect WM trace from distracting information. Although a large body of experimental evidence links neural oscillations to WM functions, theoretical understanding of their interrelations is still incomplete.

In this study, we used a computational approach to explore a potential role of beta and alpha oscillations in control of WM stability. First, we examined a single bistable module that served as a discrete object representation and was resonant in the beta-band in the active state. We demonstrated that beta-band input produced differentially stronger excitatory effect on the module in the active state compared to the background state, while this difference decreased with the input frequency. We then considered a system of two competing modules, selective for a stimulus and for a distractor, respectively. We simulated a task, in which a stimulus was loaded into the first module, then an identical oscillatory input to both modules was turned on, after which a distractor was presented to the second module. We showed that beta-band input prevented loading of high-amplitude distractors and erasure of the stimulus from WM. On the contrary, alpha-band input promoted loading of low-amplitude distractors and the stimulus erasure.

In summary, we demonstrated that stability of WM trace could be controlled by global oscillatory input in a frequency-dependent manner via controlling the level of competition between stimulus-encoding and distractor-encoding circuits. Such control is possible due to difference in the resonant and non-linear properties between the background and the active states.

## Introduction

Working memory (WM) is the process by which brain circuits hold on-line information that is necessary for multiple functions but not currently available from the sensory systems (Baddeley, 2003). Retention of information in WM is believed to be supported by sustained neural activity characterized by dynamic increase of firing rates in neural populations selective for this information (e.g. Fuster, Alexander, 1971; Goldman-Rakic, 1995; Constantinidis, Goldman-Rakic, 2002; Spaak et al., 2017; Murray et al., 2017). While multiple brain regions show WM-related persistent activity, two have been found particularly key: the prefrontal cortex (PFC) and the posterior parietal cortex (PPC) (Chafee, Goldman-Rakic, 1998; Qi et al., 2010; Salazar et al., 2012). Alongside dynamic increases of firing rates, changes in collective oscillatory activity in various frequency bands were also observed during WM tasks (Siegel et al., 2009; Haegens et al., 2010; Liebe et al., 2012; Wimmer et al., 2016; Lundqvist et al., 2016; Lundqvist et al., 2018; Kornblith et al., 2016). While such oscillatory dynamics have been linked to WM performance, their specific role has not been fully clarified. Delineating which specific mechanisms mediate interaction between the WM-related oscillations and the persistent activity to control WM functions also remains an open question.

In this paper, we focus on the beta and alpha oscillations in their potential role to flexibly control WM stability in the presence of distractors. Beta oscillations are of special interest in WM studies since they are presumably related to keeping neural activity patterns invariant over time, which was formulated as the “beta status quo” hypothesis (Engel, Fries, 2010). More specifically, in WM tasks, prefrontal beta power drops during stimulus presentation (i.e. when the existing pattern should be modified) and increases during the retention period (Siegel et al., 2009; Lundqvist et al., 2016; Lundqvist et al., 2018; Kornblith et al., 2016; Wimmer et al., 2016). Importantly, this beta increase is specific to active information maintenance (Wimmer et al., 2016) at “informative” sites, containing neurons that carry information about WM content (Lundqvist et al., 2016; Lundqvist et al., 2018). Furthermore, beta activity increases with WM load at the informative sites and decreases at the non-informative sites (Lundqvist et al., 2016; Kornblith et al., 2016). These results suggest that WM-related beta increase does not merely reflect recovery from stimulus presentation but likely plays a functional role in WM retention and gating.

Alpha oscillations are believed to reflect cortical inhibition (Klimesch, 2012). Alpha activity is usually increased in posterior regions during WM retention in a load-dependent manner (Lieberg et al., 2006; Jensen et al., 2002), supposedly protecting WM content from sensory inputs. However, alpha activity is weaker in the regions that are related to WM content retention, compared to irrelevant regions (Sauseng et al., 2009; Rösner et al., 2020). This suggests that activation of WM-retaining populations is accompanied by alpha decrease in these populations.

From the theoretical point of view, active retention of WM content is classically described in terms of bistable attractor networks (e.g. Amit, Brunel, 1997). Spiking versions of the classical WM models did not address the impact of oscillations on WM dynamics, since they required the dynamics of spiking to be asynchronous (Gutkin et al 2001), although Tegner et al. (2002) showed that oscillations could be tolerated given slow NMDA-dependent excitation. At the same time, in that work and later accounts (as in Roxin, Compte, 2016; Lundqvist et al., 2010; Lundqvist et al., 2011), oscillations did not appear to play a causal mechanistic role in WM retention and gating. In fact, despite its prominence in the experimental data, the functional role of oscillatory activity has received only limited attention in the theoretical literature. Kopell et al. (2011) presented a data-driven model, in which high beta (beta 2) and gamma oscillations in PPC concatenate to produce low beta (beta 1) oscillations that support the memory trace. This mechanism does not appear to apply in the PFC, where gamma and beta2 are shown to compete, rather than concatenate during WM retention (Lundqvist et al., 2016; Lundqvist et al., 2018; Bastos et al., 2018). This suggests that there could be a complementary PFC-based mechanism of WM retention involving beta2 oscillations.

In a complementary approach, Dipoppa and Gutkin (2013) explored how external input oscillations could control WM gating and retention in a bistable non-oscillatory network of excitatory neurons. It was shown that alpha-band input terminated WM retention and prevented new stimuli from being memorized, while theta-band input also prevented memorization, but spared WM retention. Schmidt et al. (2018) explored a similar low-dimensional model, containing either a single excitatory population or several competing populations. It was demonstrated that (1) slow oscillations promoted object memorization and spontaneous switching between objects in WM, (2) beta- / low gamma-band input erased WM content, (3) high gamma-band input promoted memorization and stabilized a metastable WM trace. These results suggest that difference in responses to an oscillatory input between background and active populations could play a major role in oscillatory WM control. However, a potential role of such differential responses in the control of competition between the retained and newly received information (and, thus, in control of neural pattern robustness) was not thoroughly studied yet.

In the present study, we propose a novel mechanism for oscillatory control of WM retention stability based on oscillation-induced changes in the level of competition between the active neural populations (selective to retained information) and the background populations (selective to irrelevant information). We consider a system of two bistable excitatory-inhibitory modules with mutual inhibition; each module has a beta-band resonance in the active state, but not in the background state. We simulate an experiment, in which a discrete stimulus is loaded into the first (S) module (switching it to the active state), and then a distractor is presented to the second (D) module, while both modules receive an identical sinusoidal input. We demonstrate that beta-band input selectively entrains the active S-module and increases its mean firing rate due to the non-linearity of the neural gain functions, which leads to increased inhibition of the D-module and a better ability of the system to ignore the distractor. Oscillations of lower frequencies (including alpha) exert an additional inhibitory effect in the active state and an excitatory effect in the background state due to the induced periodic wandering between the basins of attraction of these states. As a result, alpha-band input in our model did not increase the mean firing rate of the S-module, but provided additional excitation to the D-module during the distractor presentation, promoting the distractor loading to WM accompanied by erasure of the stimulus. In summary, we demonstrate WM-trace stabilization in the presence of a distractor by a beta-band input and its destabilization by an alpha-band input.

## Results

The basic unit of our models is a neural representation of a discrete object, implemented as a bistable network containing an excitatory and an inhibitory neuronal population. Further in the text, we refer to such network as a “module”. A module has two steady states: the background state with low firing rates, and the active state with high firing rates. When a module is in the active state; the corresponding object is retained in WM. In this paper, we considered a single-module system (for the initial analysis), as well as a system of two modules with mutual inhibition. We used the latter system to model the competition between a stimulus being retained in WM and a distractor. Most of the results in this study were obtained using low-dimensional (firing rate) models. The main findings were then confirmed by simulations of the corresponding spiking networks.

Since our aim is to explore the effects of oscillatory input on stability of WM retention, the modules did not generate oscillations by themselves. Rather, they were endowed with beta-band resonance in the active state, but not in the background state – i.e. demonstrated *state-dependent resonanc*e. Oscillations were delivered to a module as an external sinusoidal signal. In the two-module system, each module received oscillations of the same amplitude and phase. First, we considered the single-module system and explored how the oscillation-induced mean firing rate shift differed between the background and the active state (as a result of the state-dependent resonance). Then we considered the two-module system and explored how the aforementioned difference leads to oscillation-induced increase or decrease of competition between the modules.

### State-dependent resonance and mean firing rate shift in a single-module system

We first explore the persistent activity dynamics and its modulation by oscillatory inputs in a single bistable module with state-dependent resonance. The system is schematically presented in Figure 1(a). The two steady states of the model are depicted together with the corresponding gain functions in Figures 1(c,d), respectively. One can see that both steady states are located in the concave parts of the gain functions, which is crucial for the effects of oscillation-induced firing rate shift that we describe further in the paper.

**Figure 1.**
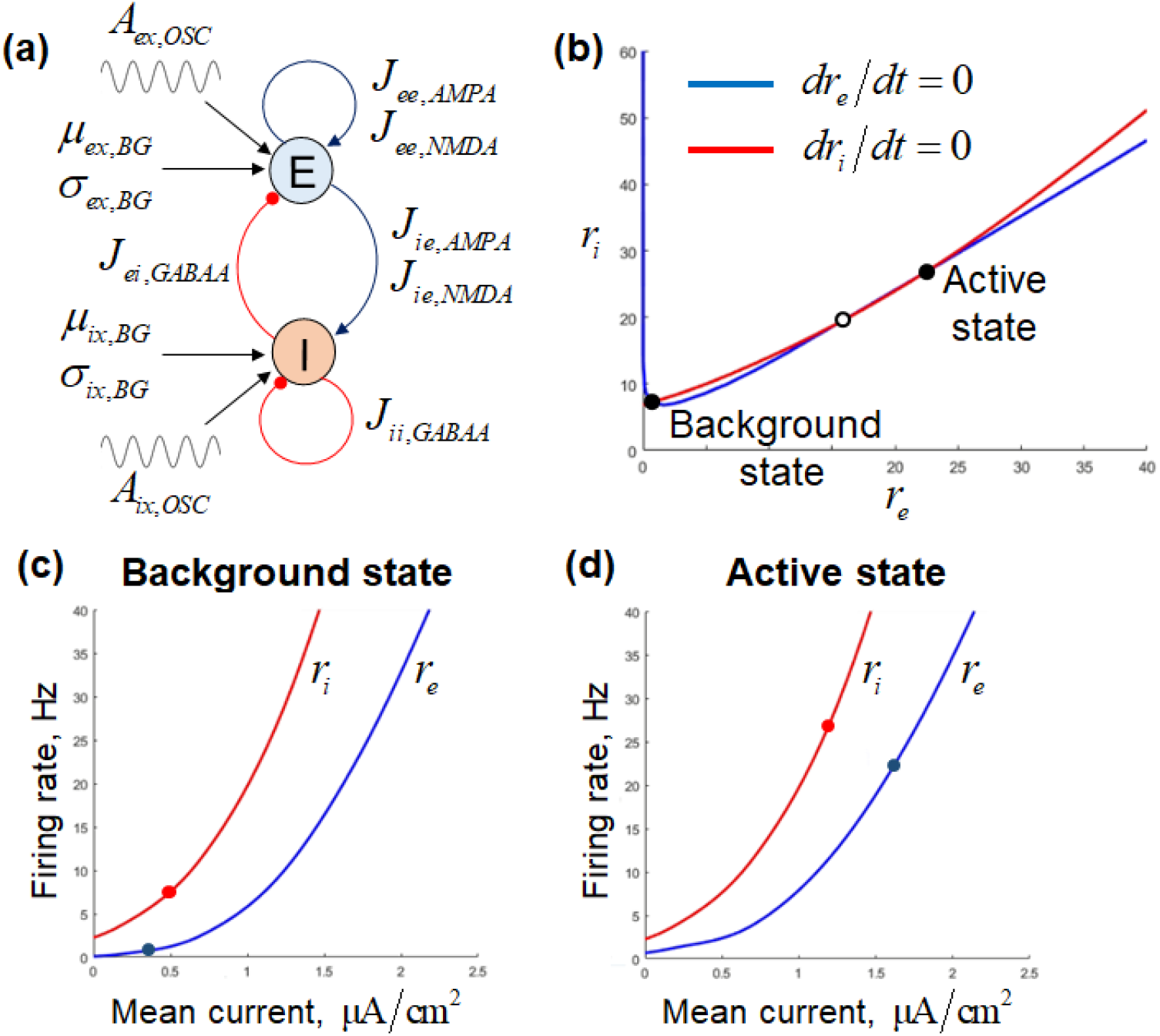
(a) Scheme of the single-module system. (b) Characteristic curves of the single-module system at the (*r_e_*, *r_i_*) phase plane. For any point on these curves, the derivatives of all the variables except of *r_e_*, *r_i_* are equal to zero. (c,d) Gain functions (firing rate versus mean input current) for a neuron in the background and the active state, respectively. Values of AMPA- and NMDA-input variances calculated at the corresponding steady-states are used both in (c) and (d).

In order to illustrate the state-dependent resonance of the model, we simulate it under oscillatory inputs in the background state and when the model is switched by a transient input to the active state (see Figure 2). Each simulation started from the background state; in Figures 2(b,d,f,h), a stimulus was presented, switching the system to the active state. In Figures 2(c – h), sinusoidal input oscillations were delivered to the system. Visually inspecting the two cases, (the background state in Figures 2(a,c,e,g)) and the stimulus-evoked active state (Figures 2(b,d,f,h), one can see that the oscillatory entrainment was much stronger when the system was in the active state. Notably, for the chosen set of parameters, the strongest entrainment was observed for the beta-band input (Figure 2(f)). Importantly, the beta-band input in the active state also provided the largest shift of the mean (time-averaged) firing rate (Δ*r_e_* = 7.4 Hz). The shifts observed in the active state for the other frequency bands (Figures 2(d,h)) and in the background state (for any frequency band) (Figures 2(c,e,g)) were considerably smaller (Δ*r_e_* = 0.5 − 2.4 Hz).

**Figure 2.**
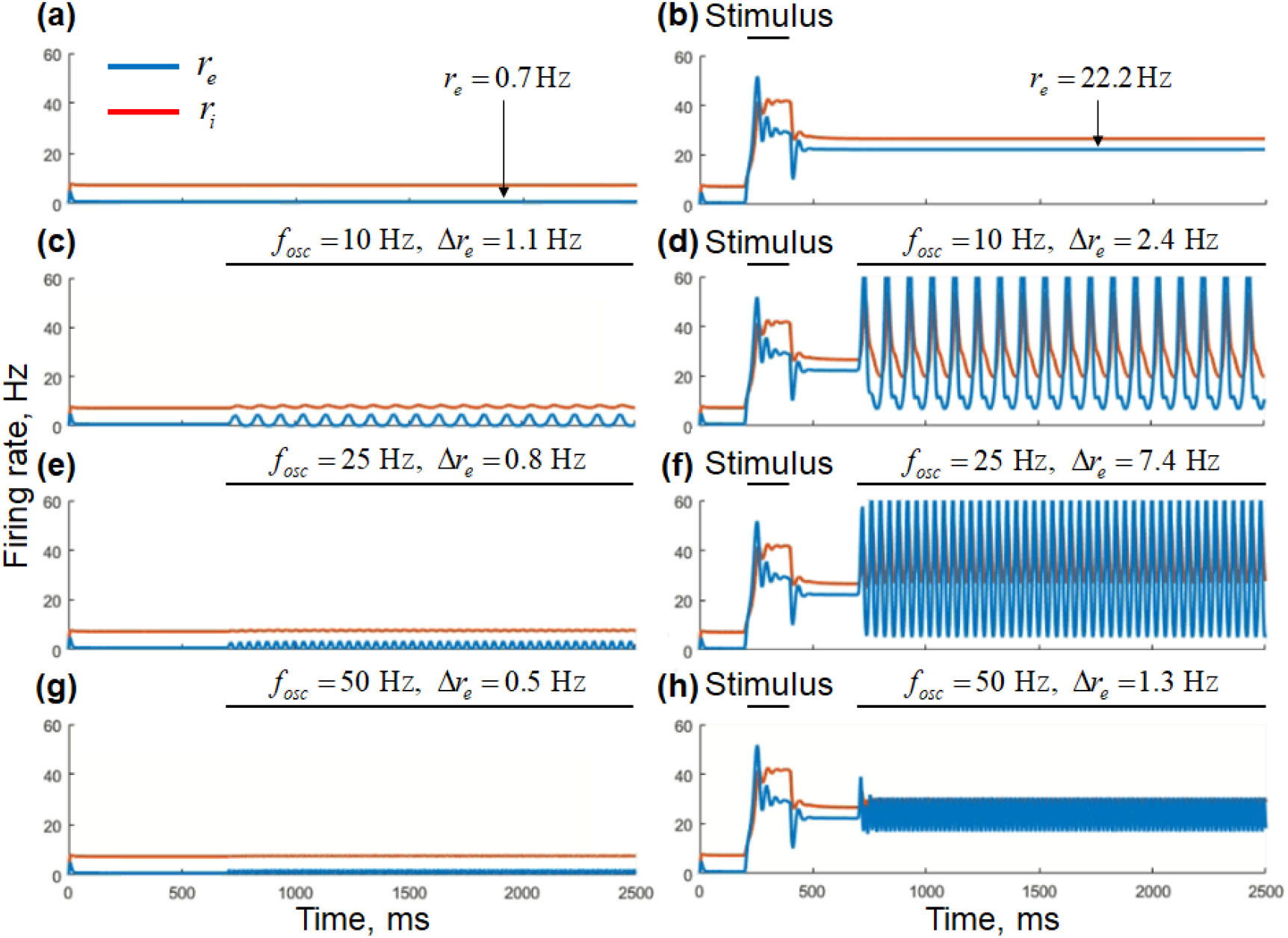
Results of the low-dimensional single-module system simulations. (a,b) – no oscillatory input; (c,d) – alpha-band input; (e,f) – beta-band input; (g,h) – gamma-band input. (a,c,e,g) – background state; (b,d,f,h) – active state. Blue curves – excitatory firing rates, red curves – inhibitory firing rates. Intervals of the stimulus and the oscillatory input presentation are marked by horizontal black lines. Δ*r_e_* time-averaged firing rate shift relative to the case without oscillatory input.

State-dependent resonance of the system can be quantified by frequency response curves (Figure 3) – amplitudes of the forced oscillations, as well as oscillation-induced shifts of the mean firing rate, vs. input frequency.

**Figure 3.**
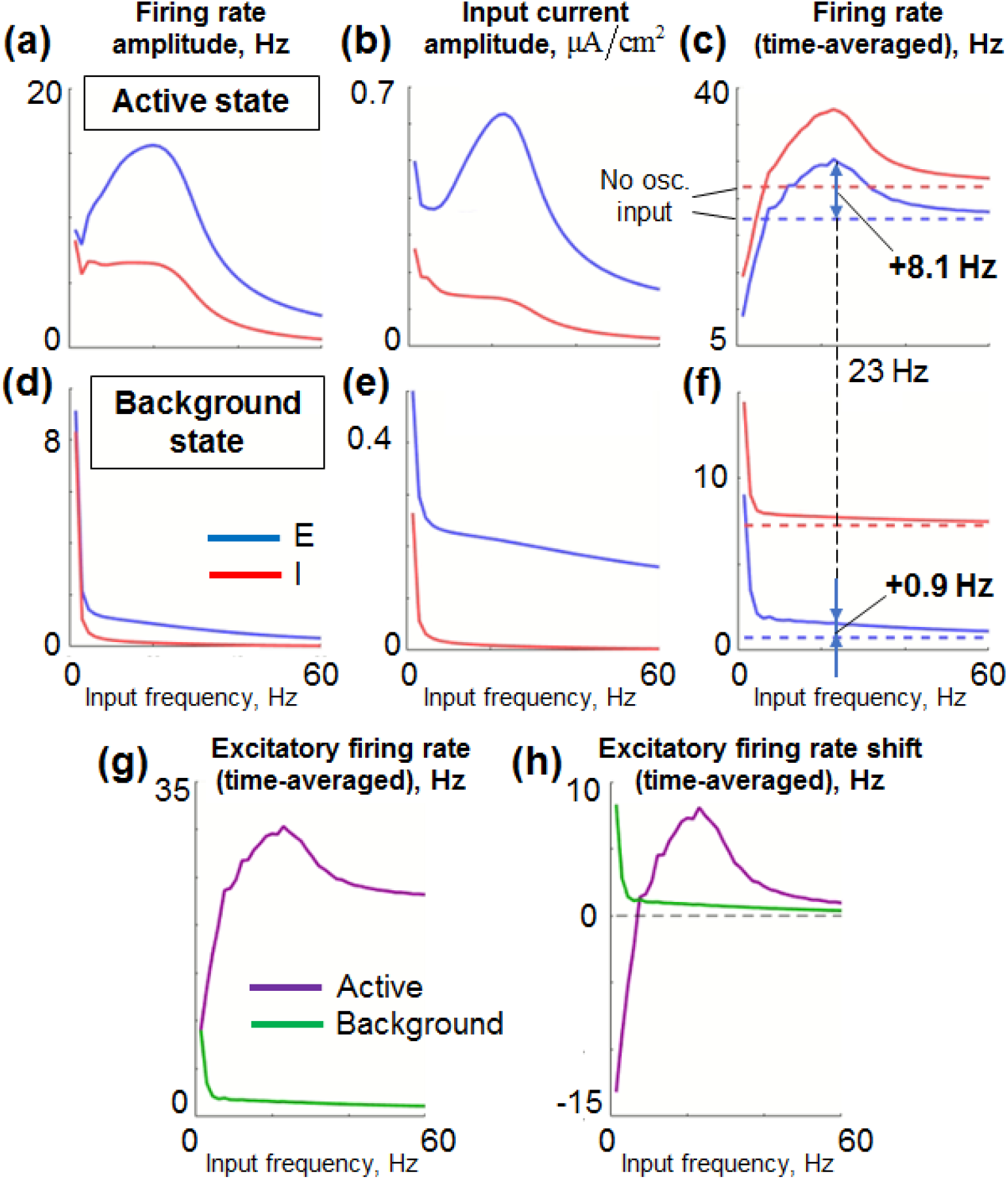
Frequency response of the low-dimensional single-module system. All graphs are functions of the input frequency. (a,d) – Firing rate amplitudes; (b,e) – total input current amplitudes; (c,f) – time-averaged firing rates. (a-c) – active state; (d-f) – background state. Blue curves in (a – f) – excitatory population, red curves – inhibitory population. Horizontal dashed lines in (c,f) represent the firing rates observed in the absence of oscillatory input. (g) – time-averaged excitatory firing rates for both states, visualized at the same plot; (h) – oscillation-induced shifts of the time-averaged firing rates (relative to the case of no input oscillations); green curves – background state, purple cures – active state. All simulations were performed for *T_sim_* = 1500 ms, and the presented quantities were calculated in the interval 750 – 1500 ms.

### Beta

Forced amplitudes demonstrate a beta-band resonant peak in the active state (Figure 3(a,b)), but not in the background state (Figure 3(d,e)). Due to the gain functions’ non-linearity (Figure 1(d)), the resonance of the amplitude leads to a resonance of the time-averaged firing rate (Figure 3(c)). Importantly, the time-averaged firing rate shift induced by the oscillatory input at the resonant frequency (~23 Hz) was much stronger for the active state than for the background state (8.1 Hz vs. 0.9 Hz; Figures 3(c,f)). The range of frequencies, in which an active population receives stronger mean excitation than a background population, could be seen in Figure 3(h) as the interval, in which the purple curve goes above the green curve. Oscillations in this range are expected to stabilize a WM trace in a multi-population system with competition by increasing the mean firing rate difference between the active and the background populations.

We can provide a heuristic explanation for the state-dependent difference in the entrainment and the mean firing rate shifts induced by the beta-band oscillatory inputs. The difference in the entrainment amplitudes between the background and the active state is related to concavity of the gain functions (Figs. 1(c,d)). Due to their concavity, higher firing rates (observed in the active state) correspond to larger derivatives of the gain functions, i.e. to stronger coupling between firing rates and synaptic currents. This, in turn, leads to stronger effective connectivity between neurons. The stronger connectivity makes a module more resonant (due to increased interaction between the excitatory and the inhibitory population), i.e. more prone to oscillatory entrainment (Ledoux, Brunel, 2011). We should note that the gain function concavity, which is crucial for the described mechanism, is a general property of neural networks operating in a subthreshold regime, typical for the neocortex (Renart et al., 2003).

The positive shift of the mean excitatory firing rate induced by the beta-band input is also related to the concavity of the gain functions. When a neuron with a concave gain function receives oscillations, its mean firing rate increases (Voronenko, Lindner, 2017). Since the oscillatory input entrains both the excitatory and the inhibitory populations of a module, both populations receive an additional mean excitatory drive (due to the non-linearity of the corresponding gain functions). We delivered the external oscillations predominantly to the excitatory populations, so the entrained amplitudes were also higher for the excitatory than for the inhibitory populations (this effect could be seen by comparing red and blue curves in Figs. 3(a,b,d,e)). The predominant entrainment of the excitatory population resulted in positive mean firing rate shifts for both the excitatory and the inhibitory populations (Figs. 3(c,f)).

### Alpha

For the lower frequencies (including the alpha band), the situation is markedly different. As the input frequency decreases, the time-averaged firing rate of the system in the active state decreases and eventually becomes even smaller than the firing rate of the unforced system (Figure 3(c)). On the contrary, the time-averaged firing rate of the system in the background state increases with decreasing input frequency; also, it is always higher than the firing rate of the unforced system (Figure 3(c)). At extremely low input frequencies, there is only one regime in the system (see the leftmost point in Figure 3(g)); thus, the system cannot serve as a WM model in this case – an effect equivalent to “WM clearance” in (Dipoppa, Gutkin, 2013.; Schmidt et al., 2018). For slightly higher frequencies, the time-averaged firing rates characterizing the active and the background regime differ, but they are close to each other (left part of Figure 3(g)). In Figure 3(h), one can see the frequency range for which the oscillatory input excites a background population stronger than the active population, or even inhibits the latter: in this range, the green line goes above the purple line. Oscillations in this range are expected to destabilize a WM trace in a multi-population competitive system and to promote updating of WM content.

We can propose a two-factor explanation for why the low-frequency oscillations have a decreased excitatory or even an inhibitory effect on a single-module system in the active state. (1) First, oscillations with a sufficiently large amplitude would force the system to wander periodically between the basins of attraction of the two steady states (background and active). During the downward phase of the entrained oscillations, the system tends towards the background state but then returns to the vicinity of the active state at the subsequent upward phase. As the frequency of the oscillations decreases, the system spends more time in the basin of attraction of the background state on each cycle of the oscillations. Consequently, the mean firing rate decreases as compared to the case of higher frequencies. (2) Second, the ratio between the oscillation amplitudes of the excitatory and the inhibitory populations gets smaller at lower frequencies (compare red and blue curves in Fig. 3(a)). Consequently, the ratio between the oscillation-induced mean excitatory drive to the excitatory and the inhibitory populations also decreases. As a result, the oscillation-induced mean firing rate shift of the excitatory population gets smaller at lower frequencies.

### Oscillatory input controls WM stability in a two-module system

In the previous section, we demonstrated that beta-band oscillations produced a stronger excitatory effect on the populations being in the active state compared to the background populations. On the contrary, oscillations of lower frequencies (including the alpha band) produced a prominent excitatory effect on the background populations, while their effect on the active populations was less excitatory or even inhibitory. These two observations served as a basis for developing a system with two active states (corresponding to retention of a stimulus and of a distractor in WM, respectively), such that switching between these states could be made harder by a beta-band input or easier by an alpha-band input.

The system contained two bistable modules with mutual inhibition between them. A schematic depiction of the system is presented in Figure 4. One module (denoted as S) served as a neural representation of a (to-be-memorized) stimulus, and the other one (denoted as D) – as a representation of a (to-be-ignored) distractor. When the stimulus or the distractor is held in WM, the S or the D module, respectively, is in the active state. Each module had the same parameters as in the single-module system described in the previous section (except of slightly increased background excitation aimed to compensate for inter-module inhibition). Inhibition between the modules was implemented by connections from the excitatory population of one module to the inhibitory population of another module. The inter-module competition was implemented using slow NMDA-based connections from the excitatory population of each module to the inhibitory population of the other module, which lead to almost tonic, non-oscillatory interaction between the modules. It was done because we were interested in the differential effect of input oscillations on both modules, and not in the oscillatory interaction between them. The strength of the mutual inhibition between the modules was set high enough, so only one of them could be in the active state at the same moment of time (i.e. the WM span was limited to one object in this model).

**Figure 4.**
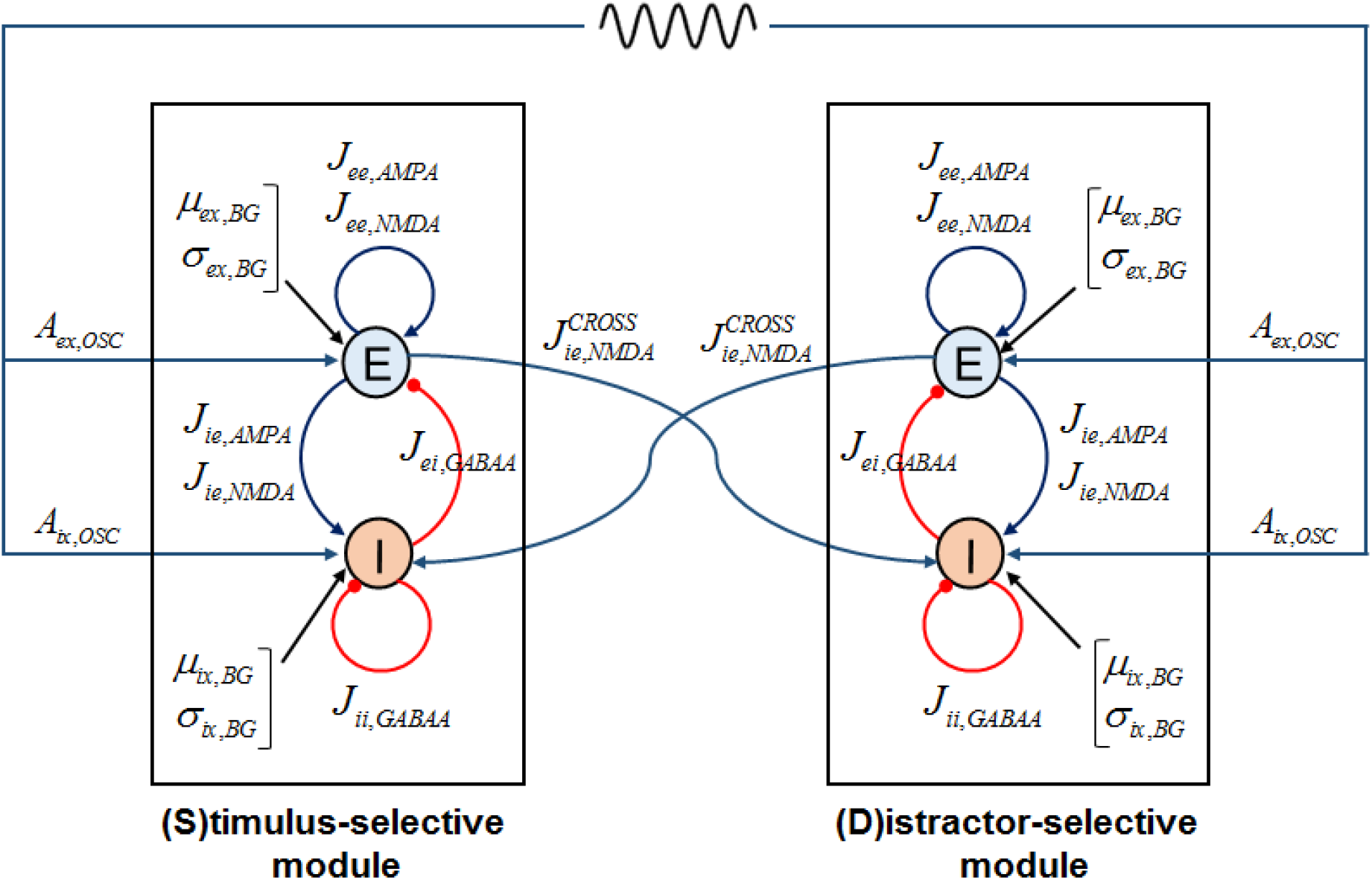
Schematic representation of the two-module system. E – excitatory populations, I – inhibitory populations.

The simulated experiment was organized as follows. First, a stimulus was presented to the S-module, switching it to the active state. Then an identical oscillatory input was presented to both modules. During the oscillatory input, the distractor was presented to the D-module. The final state of the system was assessed after the termination of the oscillatory input.

To understand how the stability of the WM trace is modulated by the oscillatory input, we analyze two versions of the numerical experiments that differ only by the distractor amplitude. In the first version, the distractor had the same amplitude as the stimulus (“strong distractor”), and its presentation to the D-module without the input oscillations was enough to switch the S-module from the active to the background state (i.e. to replace the stimulus by the distractor in WM). In the second version of the experiment, we decreased the distractor amplitude (“weak distractor”), so its presentation did not change the state of the oscillation-free system. We demonstrate that (1) beta-band input could protect the retained stimulus from being replaced by the strong distractor, and (2) that alpha-band input could promote memorization of the weak distractor with erasure of the previously retained stimulus.

### Beta-band oscillatory input protects WM from strong distractors

Simulations of the two-module system with strong distractor are presented in Figure 5. In the absence of input oscillations, the distractor replaced the stimulus in WM (Figure 5(a)). Beta-band input delivered to both modules protected the retained stimulus from the distractor presentation (Figure 5(c)). Alpha-band or gamma-band input of the same amplitude did not produce such stabilizing effect (Figures 5(b,d)).

**Figure 5.**
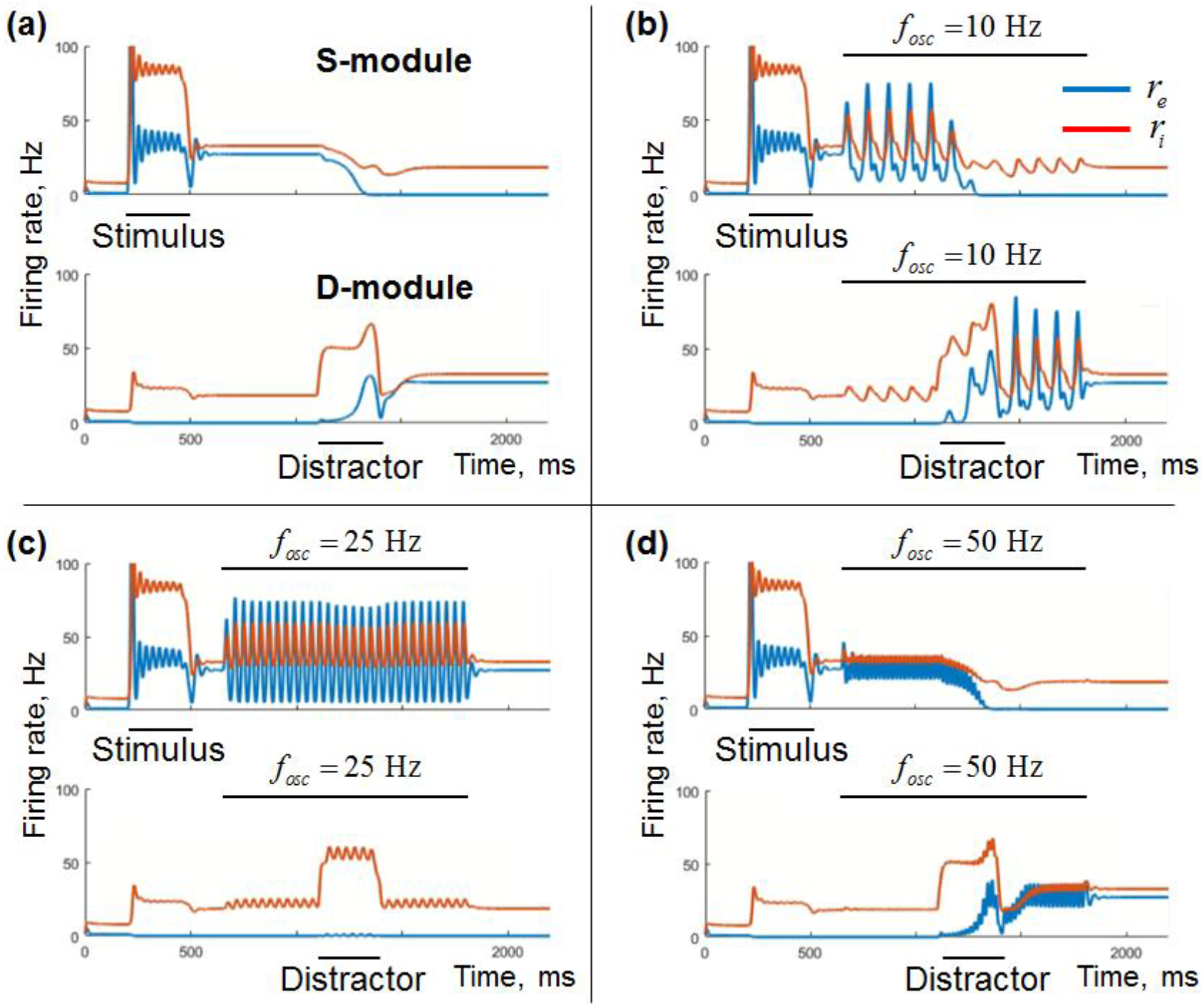
Simulation results for the low-dimensional two-module system with strong distractor. No oscillatory input, (b) alpha-band input, (c) beta-band input, (d) gamma-band input. Top subpanels – stimulus-selective module (S), bottom subpanels – distractor-selective module (D). Blue curves – excitatory firing rates, red curves – inhibitory firing rates. Intervals of stimulus / distractor presentation and of the oscillatory input are represented by horizontal black lines.

We can again explain the results by the state-dependent resonance of the system. As we saw above, each of the system’s modules demonstrated stronger beta-band oscillatory entrainment in the active state as compared to the background state. The mean excitatory firing rate shift induced by the beta-band input was also stronger in the active state. For the two-module system (4), such state-dependent resonance provided an oscillation-induced increase of competition between the modules. More specifically, a large oscillation-induced positive shift of the mean excitatory firing rate of the active (S) module produced strong inhibition of the background (D) module (via the inter-module connection), which overcame a small positive firing rate shift that beta input induces in a single background module (Fig. 5(c)). As a result, the identical beta-band input to both modules suppressed the excitatory population of the D-module, making it harder for the strong distractor to replace the stimulus in WM.

To explore in more detail how the ability of oscillations to protect WM trace from the strong distractor depends on their frequency *f_osc_* and amplitude *A_osc_*, we performed simulations for various combinations (*f_osc_*, *A_osc_*). The result is presented in Figure 6. In the region 1, the distractor was loaded into WM, erasing the stimulus. In the region 3, the oscillatory input protected the stimulus retention and blocked memorization of the distractor. The region 2 corresponds to the intermediate situation, in which the distractor was not memorized by itself, but caused erasure of the stimulus from WM. In the region 4, the background regime became unstable: even without stimulus presentation, oscillations amplified any initial asymmetry in the system and switched one of the modules to the active state (this effect is similar to the spontaneous oscillation-induced switching in a noisy multi-population system demonstrated in (Schmidt et al., 2018)). In the region 5, oscillations erased WM content and prevented stimulus memorization – the effect similar to the one described in (Dipoppa, Gutkin, 2013; Schmidt et al., 2018).

**Figure 6.**
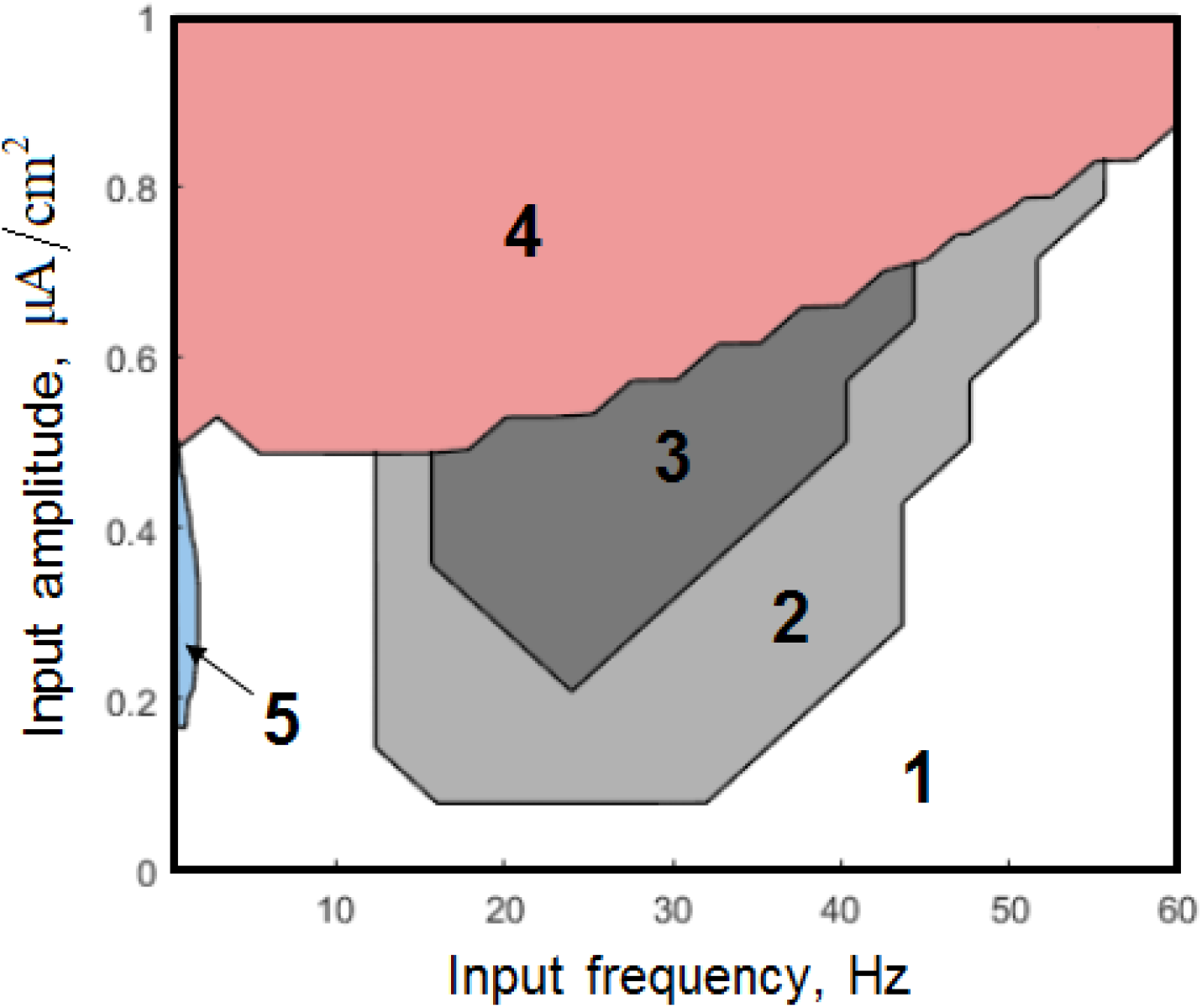
Behavior of the low-dimensional two-module system with the strong distractor for various parameters of the oscillatory input. 1 – stimulus is erased, distractor is memorized; 2 – stimulus is erased, distractor is ignored; 3 – stimulus is protected, distractor is ignored; 4 – the background regime loses stability, 5 – oscillations erase WM content and prevent stimulus loading. To find the regions 1-3, we calculated the average firing rates of the modules (S and D) after the termination of the oscillatory input (averaging window: 2000 - 2200 ms; simulation timings were the same as in Figure 5). To find the region 4, we performed a simulation without stimulus and distractor presentation and checked whether oscillations activated one of the modules (oscillatory input: 500 – 5800 ms; firing rates averaging window: 4000 – 6000 ms). To find the region 5, we performed a simulation without distractor presentation and calculated the average firing rate of the S-module (oscillatory input: 650 – 10000 ms; firing rates averaging window: 2000 – 10000 ms).

In summary, there is a range of the input oscillation amplitudes (about 0.25 – 0.5 μA/cm^2^), for which the stabilization occurred only in the frequency range limited to the beta-band. For smaller amplitudes, the oscillations were too weak to provide stabilization. For larger amplitudes, the stabilization occurred in a wider range of frequencies, including the gamma-band. Oscillations of very high amplitudes made the background regime of the system unstable.

### Alpha-band oscillatory input promotes weak distractor access to WM

Simulations of the low-dimensional two-module system with the weak distractor are presented in Figure 7. In the absence of oscillatory input, distractor presentation was ignored (Figure 7(a)). The same behavior was observed in the presence of beta-band or gamma-band input (Figure 7(c,d)). Alpha-band input destabilized stimulus retention, so the distractor was memorized, and the stimulus was erased from WM (Figure 7(b)).

**Figure 7.**
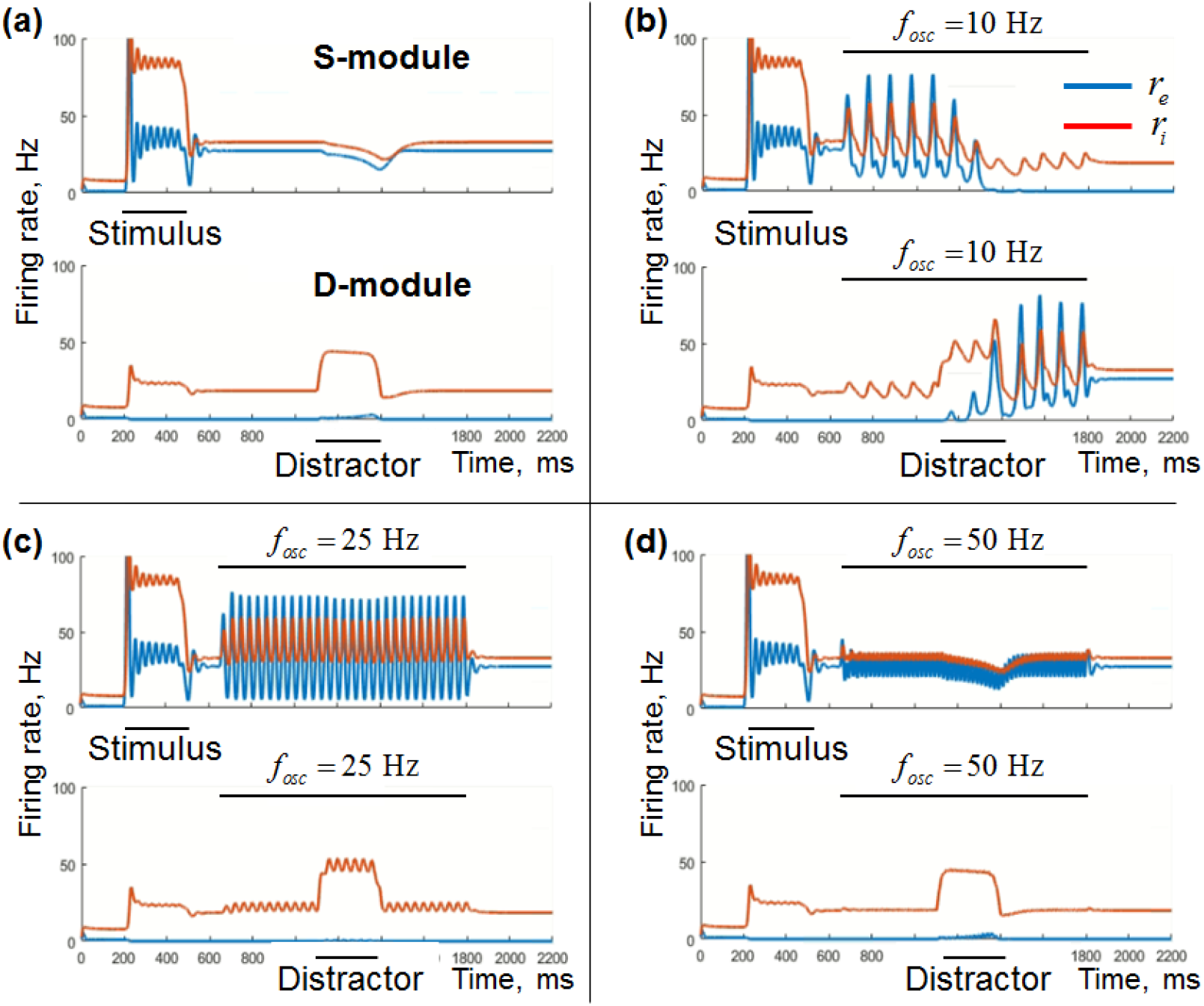
Simulation results for the low-dimensional two-module system with the weak distractor. (a) No oscillatory input, (b) alpha-band input, (c) beta-band input, (d) gamma-band input. Top subpanels – stimulus-selective module (S), bottom subpanels – distractor-selective module (D). Blue curves – excitatory firing rates, red curves – inhibitory firing rates. Intervals of stimulus / distractor presentation and of the oscillatory input are represented by horizontal black lines.

Similarly to the previous section, we explored the effect of the oscillation parameters (*f_osc_*, *A_osc_*) on their ability to promote memorization of the weak distractor. The result is presented in Figure 8 (the legend is the same as in Figure 6). For the inputs with frequencies below the beta-band (including alpha) and with high enough amplitude (0.15 – 0.5 μA/cm^2^), the distractor was memorized, and the stimulus was erased from WM (region 1). Around the region 1, there lies the region 2, in which the distractor was not memorized itself, but erased the stimulus from WM. At the frequencies roughly above 15 Hz and at low amplitudes (< 0.1 μA/cm^2^), the distractor was ignored, and the stimulus remained in the WM (region 3). As in the strong-distractor case, very strong oscillations made the background regime unstable (region 4), while moderate oscillations of very low frequency destroyed the active regime (region 5).

**Figure 8.**
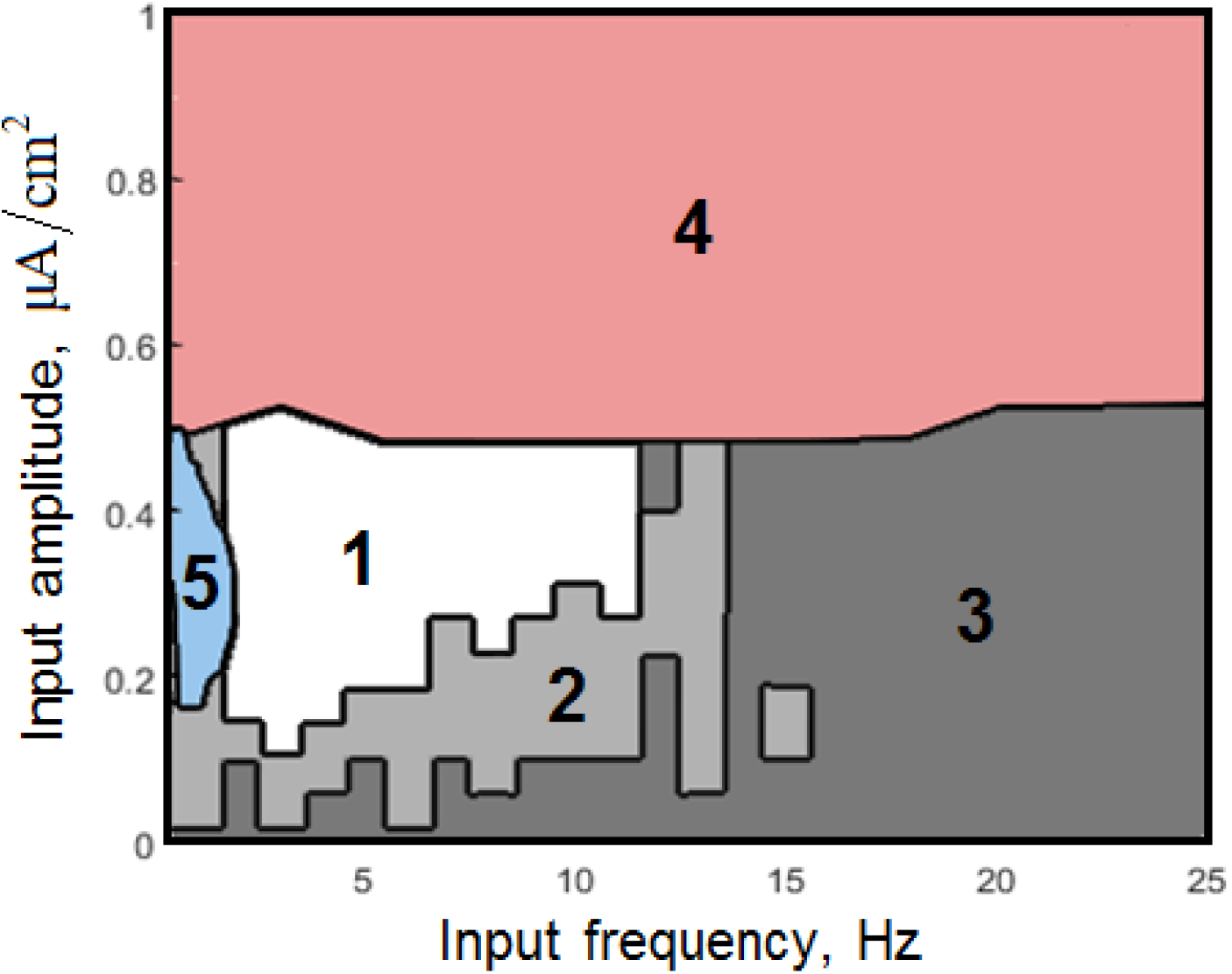
Behavior of the low-dimensional two-module system with the weak distractor for various parameters of the oscillatory input. 1 – stimulus is erased, distractor is memorized; 2 – stimulus is erased, distractor is ignored; 3 – stimulus survives, distractor is ignored; 4 – the background regime loses stability, 5 – oscillations erase WM content and prevent stimulus loading. To find the regions, we used the same procedure as in Figure 6. The borders between the regions 1, 2, and 3 are not smooth because of phase-related effects: simulation result depended on the phase of oscillations at the end of the distractor presentation and at the end of the oscillatory input itself).

The destabilizing action of the low-frequency input stems from two effects. First, as we saw in the single-module system, low-frequency oscillations produce weaker excitation (or produce inhibition) in active modules, compared to beta-band oscillations. Consequently, in the two-module system with low-frequency input (including alpha), the active S-module receives weaker oscillation-induced excitation (compared to beta input) and, thus, exerts weaker inhibition on the D-module, allowing for the distractor to initiate firing rate increase when presented. The second effect is that the D-module gets entrained in a non-linear manner by the input oscillations during the distractor presentation, which produces additional excitation to the D-module and promotes memorization of the distractor.

The described mechanism fits with the results presented in Fig. 7. Thus, in Fig. 7(b), alpha oscillations do not increase the mean firing rate of the S-module (upper panel), so the D-module (lower panel) does not receive additional inhibition, and the distractor presentation is able to initiate firing rate increase in the D-module. The distractor itself is too weak to switch the state of the system without oscillations (Fig. 7(a), lower panel). However, alpha oscillations entrain the D-module during the distractor presentation, providing an additional excitation, which is enough to switch the D-module to the active state and the S-module – to the background state (Fig. 7(b)). Thus, the alpha-band input promoted the replacement of the stimulus retained in WM by the weak distractor.

The strong mean excitatory effect of the low-frequency oscillations on the D-module during the distractor presentation is also explained by two factors. (1) First, similarly to what happens in the upper state, oscillations make the module wander between the basins of attractions of the two steady states. During the upward phase of the oscillations, the module tends towards the active state; at lower frequencies, this happens for a longer time interval, so the mean firing rate gets higher. (2) Second, the D-module is in the background state at the beginning of the distractor-related activity, so its excitatory firing rate is initially very low. Consequently, the oscillatory input induces a strongly non-linear response due to strict positivity of firing rate (Fig. 7(b), lower panel, blue curve), which, in turn, produces a strong positive mean drive to the excitatory population. This could be also understood from the fact that the gain function of the excitatory population is almost flat to the left of the background state, but rises quickly to the right of this state (see the blue curve in Fig. 1(c)).

### Dependence of WM stability on the distractor parameters

The distractor in our model was represented by a brief pulse delivered both to the excitatory and to the inhibitory population of the D-module. It was characterized by two parameters: (1) amplitude of the pulse delivered to the excitatory population (distractor strength) and (2) ratio between the amplitudes of the pulses delivered to the inhibitory and to the excitatory population (I/E ratio). In this section, we explored how these parameters interact with the oscillatory input parameters (frequency and amplitude) in defining whether the stimulus would be erased by the distractor presentation or protected by the input oscillations. We were specifically interested in the question of how fine-tuned the distractor and the oscillatory input parameters should be to provide selective stimulus protection by the beta-band oscillations.

First, we varied the distractor strength and the I/E ratio for several values of oscillations’ amplitude and probed the behavior of the system under alpha-, beta-, and gamma-band inputs. The results are presented in Figure 9. It is seen that for most of the values of I/E ratio, there exists a range of distractor amplitudes (width of the pink region), for which beta-band input selectively stabilizes the stimulus retention. This range gets larger with the increasing I/E ratio. However, as the I/E ratio gets too high (about 0.2), the stabilizing effect of oscillations becomes irrelevant, since the distractor of any amplitude is now unable to switch the state of the system (see the black regions in the top parts of the diagrams in Figure 9). In general, it is seen that the parameter region of beta-selective stabilization (i.e. the pink region) gets larger as the amplitude of oscillations increases (compare Figures 9(a), (b), and (c)).

**Figure 9.**
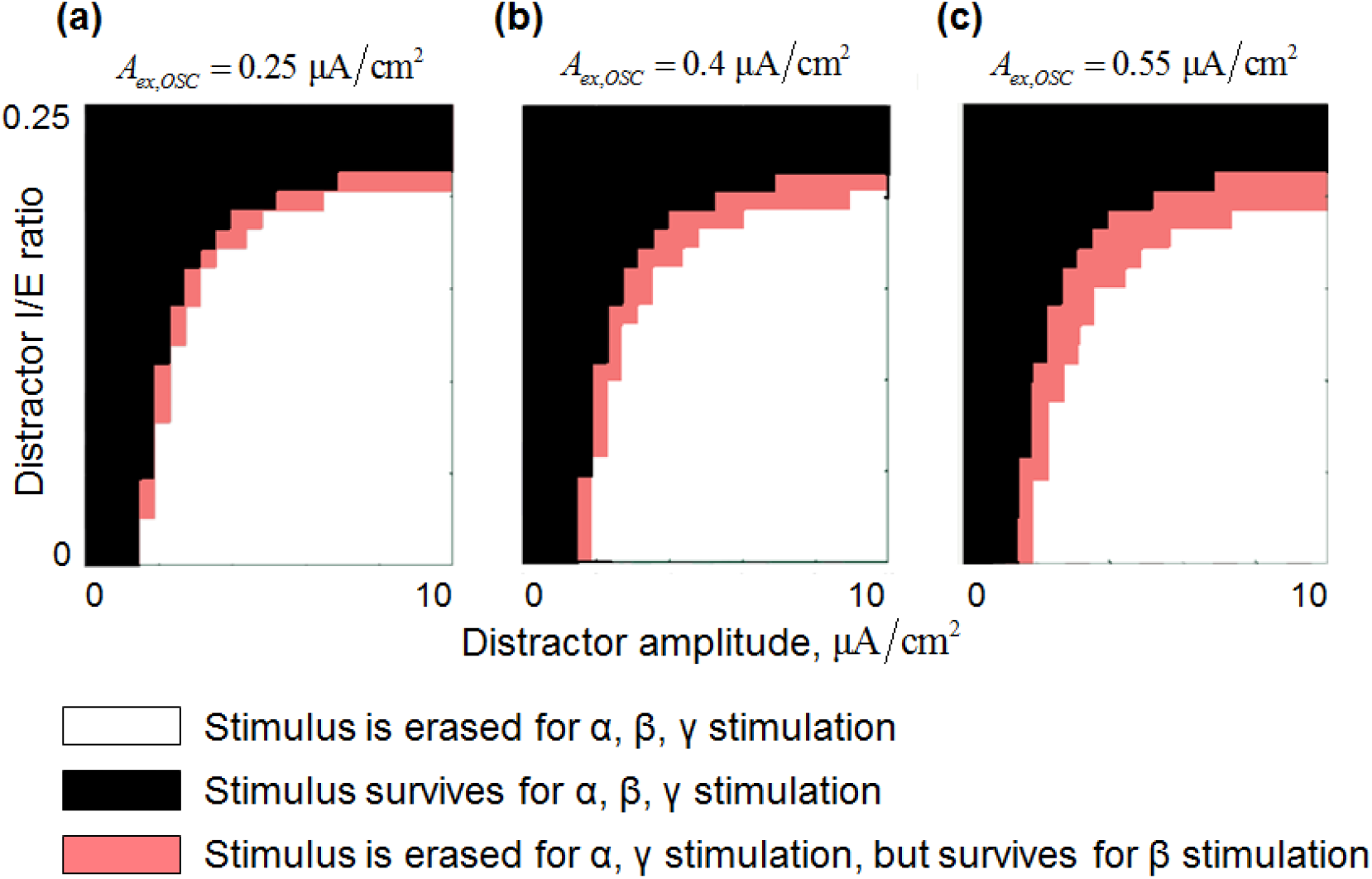
Selective stabilization of WM trace by beta-band input for various distractor parameters. Horizontal axis – distractor amplitude delivered to the excitatory population of the D-module; vertical axis – ratio between the distractor amplitudes delivered to the inhibitory and to the excitatory populations of the D-module (I/E ratio). (a,b,c) – various amplitudes of the input oscillations. Black regions – stimulus is erased by the distractor presentation during 10, 25, or 50 Hz input; white regions – stimulus survives the distractor presentation during 10, 25, or 50 Hz input; pink regions – stimulus is erased by the distractor presentation during 10 or 50 Hz input, but survives it during 25 Hz input (i.e. stimulus is selectively protected by beta oscillations). Stimulus survives (black / pink regions) when the distractor is weaker and more inhibitory. If the I/E ratio is too high, the distractor does not erase the stimulus, regardless of its amplitude (top black parts of the figures). The region of the selective beta-band stabilization (pink) is wider for stronger oscillatory input: (c) > (b) > (a). The interval of the distractor amplitudes, for which the selective beta-band stabilization occurs, is wider when the I/E ratio is higher.

Second, we varied the distractor amplitude and the frequency of the input oscillations for several values of the input oscillations’ amplitude and for two values of the distractor I/E ratio. The results are presented in Figure 10. Each curve corresponds to a certain input amplitude value. In the region above a curve, the distractor presentation switches the system, erasing the previously retained stimulus from WM; in the region below the curve, the stimulus retention is spared. One could see a stabilizing effect of beta oscillations: for the inputs in the beta band (purple regions), a larger distractor amplitude is required to erase the stimulus from WM. A destabilizing effect of lower frequencies (below beta) is also seen: for the frequencies from this band (yellow region), a smaller distractor amplitude is enough to erase the stimulus from WM. The ranges of the distractor amplitudes for which the stabilizing or the destabilizing effects occur are larger for stronger oscillations: the green curves in Figure 10 go further upwards in the purple region and further downward in the yellow region, compared to the red and the blue curves. These ranges are also larger for the higher distractor I/E ratio: the peaks in the purple regions and the troughs in the yellow region are more pronounced in Figure 10(b) compared to Figure 10(a).

**Figure 10.**
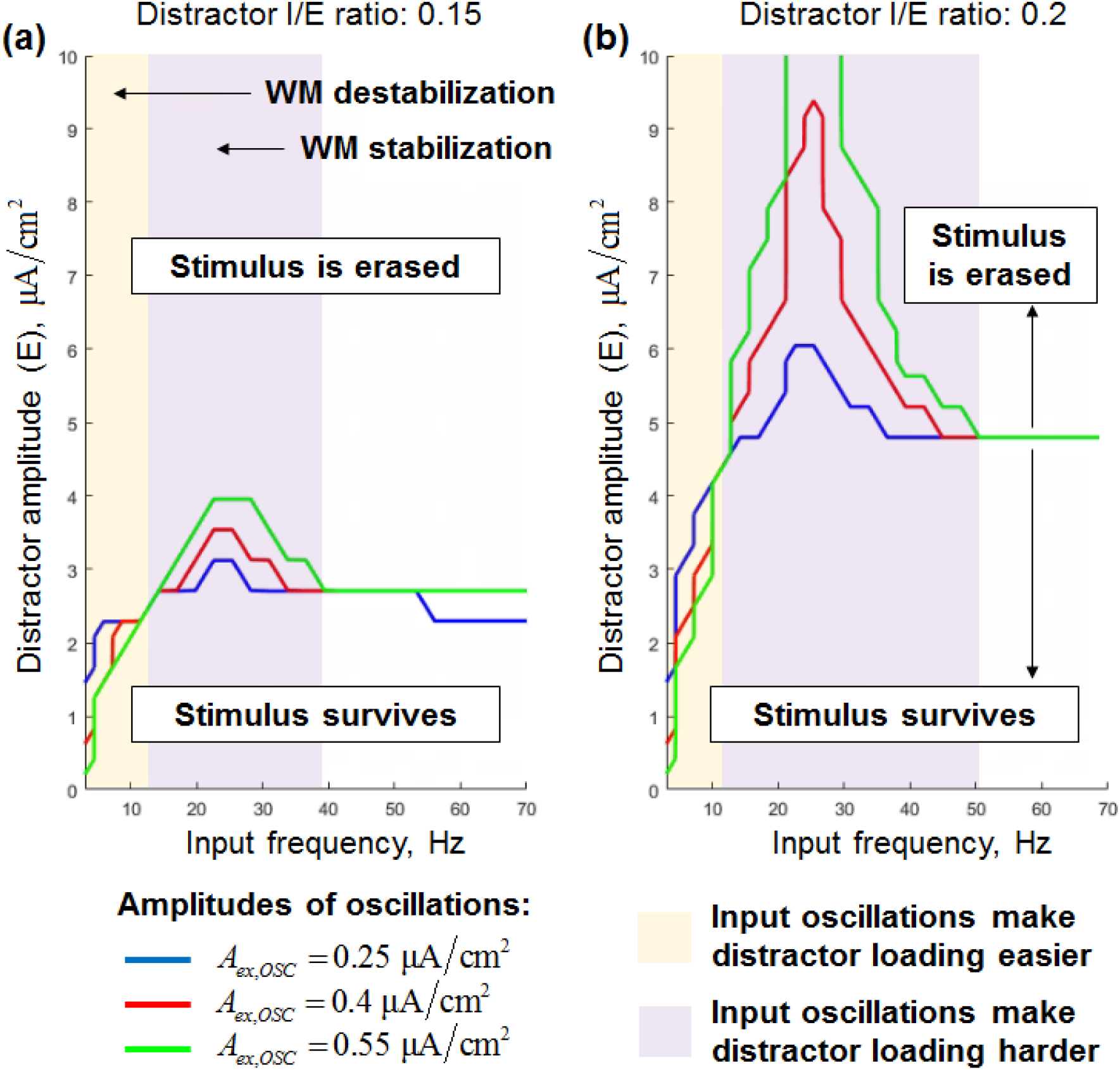
Selective stabilization of WM trace by input oscillations for various distractor and oscillations parameters. Horizontal axis – frequency of the input oscillations; vertical axis – amplitude of the distractor received by the excitatory population of the D-module. (a) and (b) correspond to two different ratios between the distractor amplitudes received by the inhibitory and by the excitatory populations of the D-module (I/E ratio). Colored curves correspond to different amplitudes of the input oscillations. For the parameter combinations above a curve, the stimulus is erased from WM by the distractor presentation; for the parameters below a curve – the stimulus retention is spared. Beta-band input stabilizes WM, making it harder for the distractor to erase the stimulus (purple region); low-frequency input (including the alpha-band) destabilizes WM, making the erasure easier (yellow region). Both effects (stabilization and destabilization) are more pronounced (observed for a larger range of the distractor amplitudes) when the distractor I/E ratio is higher and the amplitude of the oscillations is larger. The results were obtained for the stimulus amplitude of 5.5 μA/cm^2^ and the stimulus I/E ratio of 0.2.

From both analyses, we see that an increase of the distractor I/E ratio reduces the required level of fine-tuning for the distractor amplitude; however, this increase itself is limited by the value above which the distractor effect becomes too weak. At the same time, an increase of the input oscillations’ amplitude reduces the required joint fine-tuning for the distractor amplitude and the I/E ratio. The amplitude increase is limited by the level above which oscillations compromise bistability (see Figures 6 and 8). In summary, we suggest that there is enough freedom in selection of the parameters for the distractor and for the input oscillations.

### Simulations of equivalent spiking networks confirm the analysis

We confirmed the results obtained in the previous sections on the low-dimensional models by reproducing them in simulations of the equivalent spiking network models.

Single-module spiking network model simulations are presented in Figure 11. They are qualitatively similar to those obtained for the low-dimensional model (Figure 2). Importantly, the strongest oscillation-induced shift of the time-averaged firing rate was observed for the beta-band input in the active state (Figure 11(f)).

**Figure 11.**
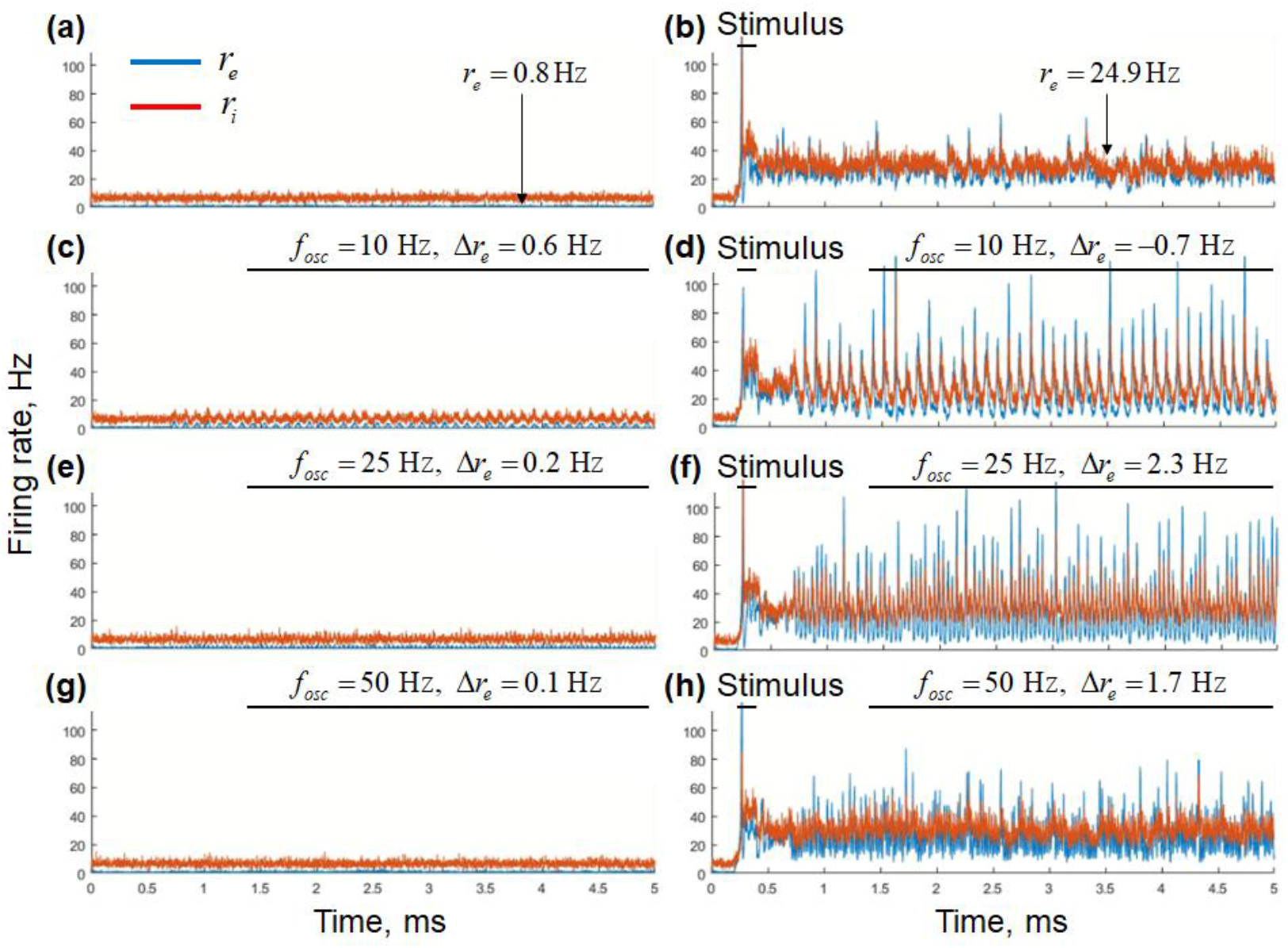
Simulation results for the spiking network model of the single-module system. (a,b) No oscillatory input; (c,d) alpha-band input; (e,f) beta-band input; (g,h) gamma-band input. (a,c,e,g) background state; (b,d,f,h) active state. Blue curves – excitatory firing rates, red curves – inhibitory firing rates. Intervals of the stimulus and the oscillatory input presentation are marked by horizontal black lines. Δ*r_e_* time-averaged firing rate shift relative to the case without oscillatory input.

The simulation results for the spiking network model of the two-module system are presented in Figure 12 for the case of the strong distractor and in Figure 13 for the case of the weak distractor. The main observations were confirmed: (1) stimulus retention was selectively protected from the strong distractor by the beta-band input (Figure 12(c)) and (2) memorization of the weak distractor was selectively promoted by the alpha-band input (13(b)).

**Figure 12.**
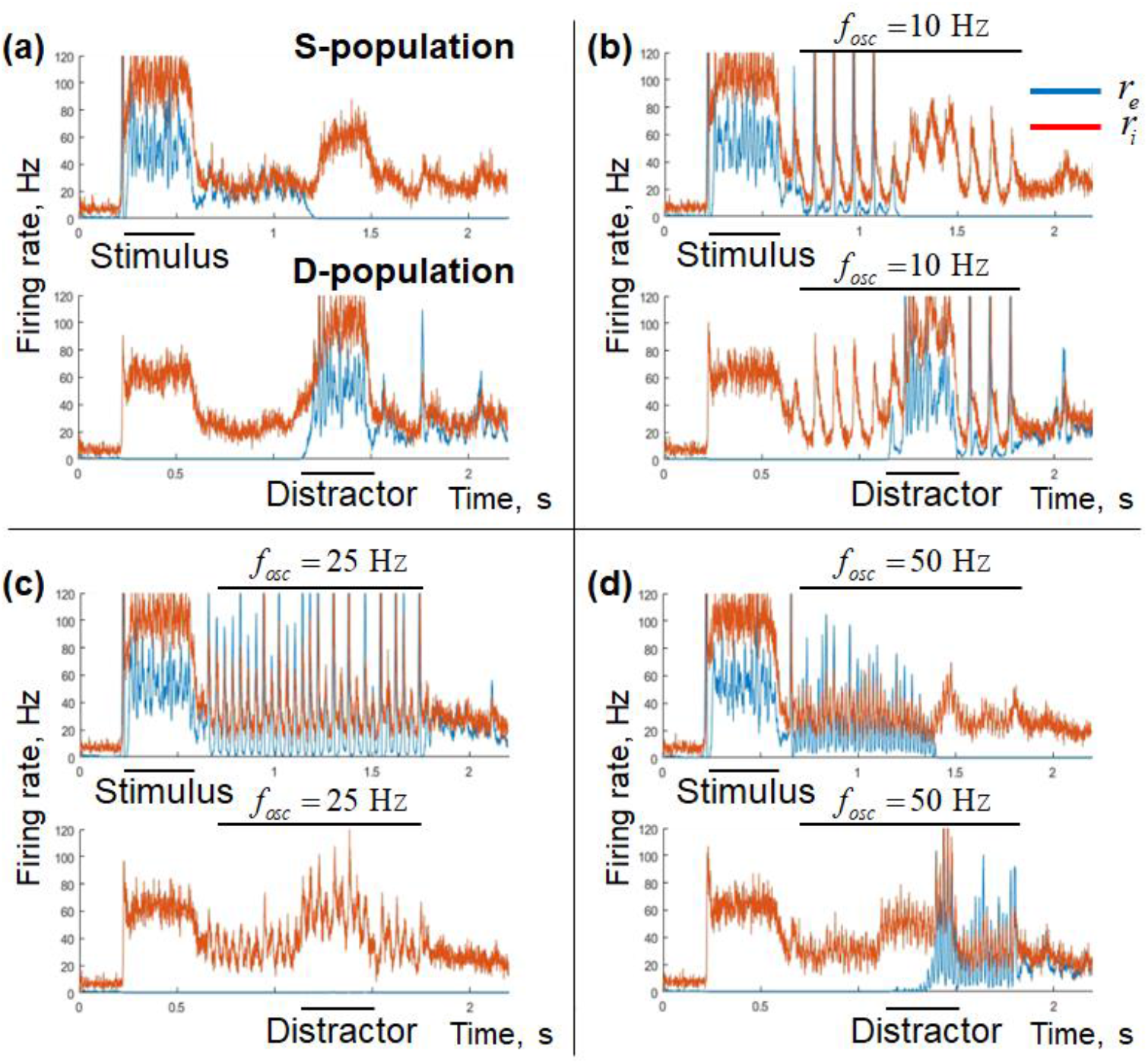
Simulation results for the spiking network model of the two-module system with the strong distractor. (a) No oscillatory input, (b) alpha-band input, (c) beta-band input, (d) gamma-band input. Top subpanels – stimulus-selective module, bottom subpanels – distractor-selective module; blue curves – excitatory firing rates, red curves – inhibitory firing rates. Intervals of stimulus / distractor presentation and of the oscillatory input are represented by horizontal black lines.

**Figure 13.**
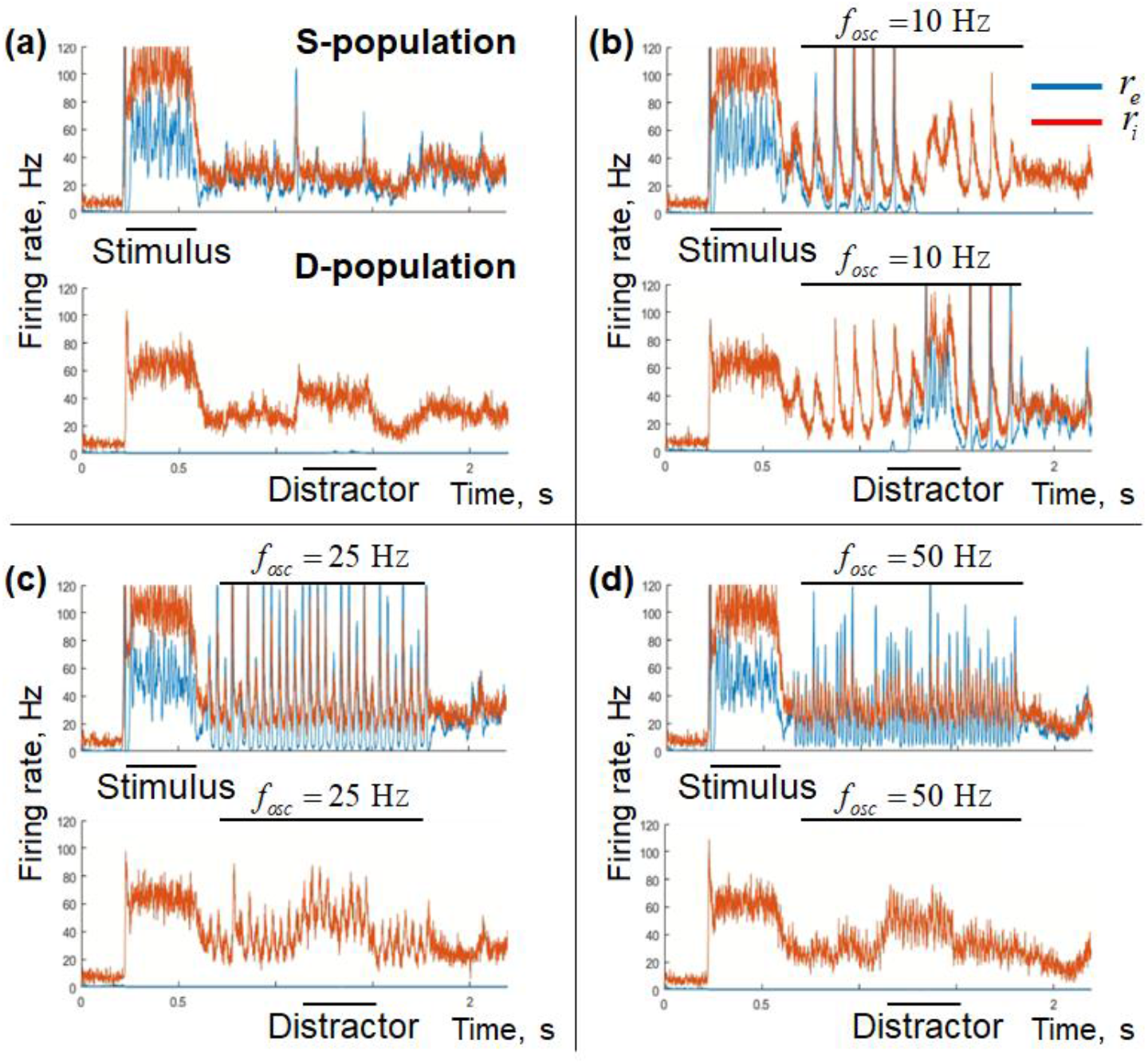
Simulation results for the spiking network model of the two-module system with the weak distractor. (a) No oscillatory input, (b) alpha-band input, (c) beta-band input, (d) gamma-band input. Top subpanels – stimulus-selective module, bottom subpanels – distractor-selective module; blue curves – excitatory firing rates, red curves – inhibitory firing rates. Intervals of stimulus / distractor presentation and of the oscillatory input are represented by horizontal black lines.

Dependence of the two-module spiking network behavior on the oscillatory input parameters is presented in Figure 14. The main findings obtained in the previous sections were confirmed. (1) For the strong-distractor case, stimulation in the intermediate amplitude values provides WM stabilization with high probability when the frequency of stimulation is in the beta band. (2) For the weak-distractor case, memorization of the distractor was promoted with high probability when stimulation amplitude was high enough and its frequency was low (below the beta-band, including the alpha band).

**Figure 14.**
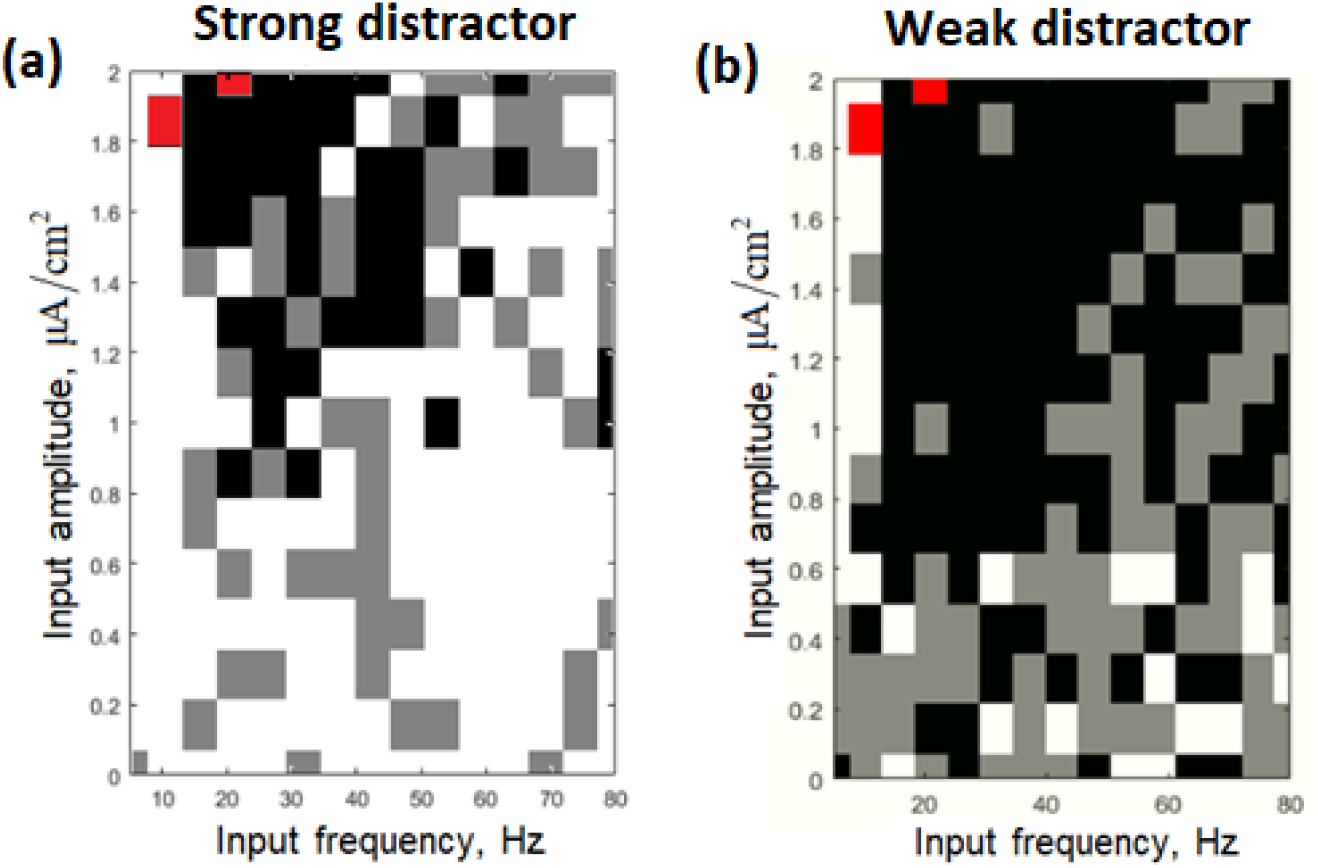
Behavior of the spiking network model of the two-module system for various parameters of the oscillatory input. The intensity of the black color represents the percentage of simulation runs in which the stimulus-related activity survived the distractor presentation (white – 0%, black – 100%; 4 runs were performed in total for the strong distractor simulations and 2 runs – for the weak distractor simulations). The stimulus-related activity level was assessed as the average over 2000 – 2200 ms window (simulation timings were the same as in Figures 12 and 13). For the parameter combinations marked by the red color, stability of the background regime was destroyed by the oscillations (assessed in a single run). To determine the red region, we performed a simulation without stimulus and distractor presentation and checked whether oscillations activated one of the modules (oscillatory input: 650 – 1400 ms; firing rates averaging window: 1100 – 1400 ms).

## Discussion

In the present study, we showed theoretically that WM stability / robustness to distractors can be controlled by externally applied oscillations. We proposed a mechanism for such control based on the state-dependent resonance acting in conjunction with the gain function non-linearities. According to this mechanism, a global beta-band input preferentially entrains WM-encoding (active) modules and increases their mean firing rate (compared to background modules). Inputs of lower frequencies (below beta) provide relatively less excitation to the active modules and more excitation to the background modules. To explore the consequences of these effects on stimulus-distractor interactions, we considered a system of two competing modules, selective for a stimulus and for a distractor, respectively. Using this system, we simulated a task, in which a stimulus was loaded into WM, then global oscillatory input was turned on, after which a distractor was presented. We demonstrated that beta-band input protects the WM trace from a competing distractor, while alpha-band input could promote replacement of WM content by the distracting information. We also demonstrated that input at very low frequency could prevent the system from retaining any WM content (similar the results were obtained for alpha oscillations in (Dipoppa, Gutkin, 2013) and for beta-gamma input in (Schmidt et al., 2018)). Importantly, the same oscillatory signal was delivered both to the WM-encoding module and to the distractor-selective module. Thus, the oscillations served as a global controller of robustness vs. flexibility trade-off, defining the system’s ability to perform WM functions, as well as its predisposition towards retaining or updating the current WM content.

The stabilizing effect of the beta-band input in our model stems from much stronger oscillatory entrainment of the stimulus-selective module (which is in the active state after the stimulus presentation), compared to the distractor-selective module (which is in the background state). Strong entrained oscillations of the stimulus-selective module result in a positive mean firing rate shift in its excitatory population. This shift leads to an additional inhibition of the distractor-selective module due to the inter-module excitatory-to-inhibitory connections. This effectively increases the cross-competition between the two modules. As a result, a distractor that is strong enough to disrupt the existing WM trace retention without oscillations, is unable to do so in the presence of the beta-band input.

Low-frequency oscillations (below the beta band) do not produce strong excitation (or produce inhibition) in the active stimulus-selective module, so the background distractor-selective module receives weaker inter-module inhibition. In such conditions, presentation of a distractor initiates activity increase in the distractor-selective module, which is additionally amplified by non-linear oscillatory entrainment. These effects are explained by (1) oscillation-induced wandering between the basins of attraction of the two steady states, which causes an average inhibitory effect in the active module and an excitatory effect in the background module, (2) entrainment of the inhibitory population of the active module, and (3) rectified character of the forced oscillations near the background state due to saturated neural gain functions at low firing rates. As a result, WM retention is destabilized, since a distractor becomes able to switch the state of the system and replace the stimulus in WM, even if it is too weak to make it without oscillatory input.

In sum total, state-dependent resonance and non-linear properties of the WM modules allow a global beta-band input to solidify the WM trace (protecting it from distractors) and a similar low-frequency (e.g. alpha-band) input – to make the WM system more flexible and labile to newly incoming stimuli. We first modeled and analyzed this oscillatory control using a low-dimensional firing rate model of two-item discrete working memory and then verified the results by direct simulations of an equivalent spiking network model. In addition, we partially explored the parameter space and demonstrated that the stabilizing / destabilizing effects do not require strong fine-tuning of the parameters.

### Relationship to experimental data

#### Beta oscillations

Stabilization of WM trace by the beta-band activity in our model is based on **two principles**: (I) **state-dependent resonance**(i.e. stronger entrainment of active populations by the beta oscillations compared to background populations) and (II) **oscillation-induced shift of the mean firing rate**.

(I)

State-dependent beta-band resonance could potentially explain several experimental findings that concern delay-period beta-band activity. (1) Beta activity increases in a load-dependent manner at “informative” sites, i.e. at the sites that either have WM-selective neurons in the closest vicinity (Lundqvist et al., 2016) or are located contralaterally to the retained stimuli (Kornblith et al., 2016). (2) Beta activity itself may contain information about WM content (Wimmer et al., 2016). (3) More information could be decoded from spiking activity at the specific phases of beta LFP (Siegel et al., 2009).

The load-dependent beta increase at the “informative” sites could be explained by our model in the following way. It is likely that the environment of an “informative” site contains many neurons from the populations selective for certain features of possible WM content. When a certain object is retained in WM, the populations representing this object are in the active state and, according to our hypothesis, they get strongly entrained by the ongoing beta oscillations. Consequently, these populations contribute to the beta-band LFP signals recorded at the nearby sites. With increasing WM load, more WM-selective populations are in the active state, so the overall level of the beta-band activity produced by the state-dependent entrainment also increases (which was observed in (Lundqvist et al., 2016; Kornblith et al., 2016)).

Load-dependent beta increase would be detected at individual sites if populations selective for different features are spatially intermingled (in this case, several representations could contribute to the LFP signal at the same site). Furthermore, if the surroundings of a site contain non-equal numbers of neurons selective for different objects, then the beta-band LFP activity at this site (generated by the beta-entrained active populations) would depend on the identity of the retained object. In other words, this beta LFP signal would carry information about the WM content, like in the results by Wimmer et al. (2016).

Our model could also explain the fact that the amount of information about the WM content is higher at the specific phase of beta LFP (Siegel). When a stimulus is retained in WM, this amount increases with the firing rate difference between the populations that are selective to the retained stimulus (which are in the active state) and other populations (which are in the background state). Our model predicts that the firing rate of an active population oscillates in the beta band with much higher amplitude than the firing rate of a background population. Consequently, the firing rate difference between them would also oscillate in the beta band, and so will do the amount of information about the WM content. As a result, this amount would be the largest at the LFP phase in which the firing rate of the stimulus-selective population reaches its maximum.

(II)

Experimental support for our second principle, i.e. oscillation-induced mean firing rate shift, is less conclusive. According to our model, beta activity provides an additional excitation to the WM-encoding populations, thus increasing the amount of information about the WM content present in the system. On the one hand, the temporal profile of the firing rate for the delay-selective cells (i.e. the cells that encode the WM content during the delay) is similar to the temporal profile of the beta-band LFP activity (Lundqvist et al., 2016). On the other hand, it was shown that the beta-band activity occurs as a sequence of brief bursts, and the amounts of information about the WM content carried by the spikes inside and outside these bursts (calculated on the whole set of recorded neurons) do not differ significantly from each other during the delay period (Lundqvist et al., 2018).

Despite the aforementioned controversy, we predict there should be a subset of the delay cells that increase their firing rates in a stimulus-selective manner during beta bursts and thus behave in agreement with our model. The existence of these cells does not contradict the absence of group-level effect in Lundqvist et al. (2018) since different cells may have opposite relations between their firing rate and beta. If such subset of neurons is found, it would be useful to explore their possible role in WM stabilization – by analyzing their activity during and after the distractor presentation, as well as during the delay period in erroneous trials. We also suggest that it would be reasonable to perform such investigation using longer delay periods since the effects related to the beta increase in WM tasks (such as the load dependence and the relation to behavioral performance) usually manifest at later delay times (Kornblith et al., 2016; Zavala et al., 2017).

#### Alpha oscillations

According to our model, we would expect alpha decrease in WM-encoding populations during the retention, since the WM content should not be replaced by distracting or irrelevant information. We suggest that the existing experimental data partly corresponds to this prediction; however, its interpretation is not straightforward.

Most of the data concerning alpha activity in WM tasks relates to the posterior brain regions. In many WM tasks, posterior alpha changes similarly to the prefrontal beta: it decreases during stimulus presentation and increases during the retention interval (Jokisch, Jensen, 2007), with this increase being stronger under higher WM load (Lieberg et al., 2006; Jensen et al., 2002). Furthermore, the posterior alpha activity is stronger before distractors than before to-be-memorized stimuli (Freunberger et al., 2009; Bonnefond, Jensen, 2012). The common interpretation of the posterior alpha increase is that it provides protection of WM content from incoming sensory information. The similarity between the prefrontal beta and the posterior alpha led to a hypothesis that these oscillations represent the same functional phenomenon with varying frequencies across different brain regions (Lundqvist et al., 2020). From this perspective, alpha and beta oscillations should have similar effects on the population activity, which conflicts with our predictions.

Despite the similarities, however, there is a notable difference between the prefrontal beta and the posterior alpha in WM tasks: the delay-period beta is weaker at the task-irrelevant sites compared to the relevant ones (Lundqvist 2016; Kornblith, 2016), while this relation is opposite for the alpha (Sauseng et al., 2009; Rösner e al., 2020; Haegens et al., 2010). As an example, the delay-period alpha activity contralateral to the memorized items decreased with the number of these items, while the alpha activity contralateral to the distractors increased with their number (Sauseng et al., 2009). A similar effect was observed in a paradigm with retro-cueing: alpha decreased contralaterally and increased ipsilaterally to the hemifield cued as relevant (Rösner et al., 2020). From these findings, one could suggest that the posterior alpha serves a protective role in WM retention, but it presumably comes, in a large part, not from those circuits that participate in WM retention, but from their interfering surroundings. It is possible that the circuits involved in a specific WM task constitute a relatively small part of the posterior network (in contrast to the PFC, where the circuits are able to flexibly change their tuning in a task-dependent manner); this latter assumption may explain the overall posterior alpha increase during the retention. Thus, we could speculate that alpha activity in WM-related circuits decreases during the retention, although this decrease is masked by a dominating increase of alpha activity in the task-irrelevant part of the network.

Several studies demonstrated an alpha decrease in WM tasks. Some of them (Gevins et al., 1997; Scharinger et al., 2017) required cognitive manipulation or did not have a well-defined retention period (e.g. if the n-back task was used). The alpha decrease in such studies could be attributed to an overall increase in the level of attention and arousal, in parallel to WM retention. More importantly, even in the case of the overall posterior alpha increase, different parts of the posterior region could demonstrate the opposite load effects: alpha increases with WM load in the occipital cortex (lower-level) but decreases with WM load in the medial parietal cortex (higher-level; Michels et al., 2010). Furthermore, a careful temporally and spatially resolved analysis could reveal multiple cortical regions (including frontal areas) with load-dependent alpha decrease (Proskovec et al., 2019). These findings further support the hypothesis that alpha decrease may reflect engagement of neural populations in WM retention process.

Since alpha activity is, in large part, a correlate of neural inhibition (Klimesch, 2012), the alpha decrease in WM-related populations could be simply interpreted as functional disinhibition of these populations when their activity is required for WM retention. Our model predicts that the alpha decrease additionally leads to more robust WM retention in the face of distracting neural activity. This prediction does not contradict the disinhibition hypothesis; however, it still requires direct experimental confirmation.

### Relationship to other models

There are multiple modeling studies that consider oscillations in WM tasks (Lisman Idiart, 1995; Tegner et al., 2002; Ardid et al., 2010; Lundqvist et al., 2010; Kopell et al., 2011; Lundqvist et al., 2011; Chik, 2013; Dipoppa, Gutkin, 2013; Roxin, Compte, 2016; Fiebig, Lansner, 2017; Pina et al., 2018; Schmidt et al., 2018; Sherfey et al., 2020). Among them, there are two studies dedicated to external oscillatory control of WM functions that are closely related to our work (Dipoppa, Gutkin, 2013; Schmidt et al., 2018).

The main commonality between our model and the models described in (Dipoppa, Gutkin, 2013; Schmidt et al., 2018) is the state-dependent resonance (strong resonant properties in the active state, but not in the background state); however, the underlying mechanisms are different. The models by (Dipoppa, Gutkin, 2013; Schmidt et al., 2018) contain only excitatory populations with neurons being in the supra-threshold regime (i.e. regularly spiking). The resonance in these models stems from spike-to-spike synchronization, and the resonant properties of each state depend on its mean firing rate. In the lower state, the firing rate is not enough to support resonant behavior, while in the active state the resonant frequency is close to the mean firing rate. In our model, the resonance stems from the interaction between excitatory and inhibitory populations, and the resonant properties of each state depend on the strength of synaptic coupling. The model operates in the subthreshold regime, so the neural gain functions are concave, and their slope is higher in the active state, which leads to increased effective coupling and a prominent resonant peak. The subthreshold spiking regime is in agreement with the experimental data, which shows high CV values during WM retention (Compte et al., 2003).

All three models rely on external oscillatory input. The input is sinusoidal in our model and pulse-like in (Dipoppa, Gutkin, 2013; Schmidt et al., 2018). We suggest that sinusoidal input is more realistic than pulse-like input since oscillations in PFC were reported to come in narrow-band bursts during WM tasks (Lundqvist et al., 2016).

Most importantly, we made a step further from (Dipoppa, Gutkin, 2013; Schmidt et al., 2018) and explored the ability of oscillations to block or promote distractor loading in a two-module competitive system during stimulus retention. The aforementioned studies did not pose this question directly and were mainly focused on single-module systems. However, we could note two underlying effects that are common for our study and the previous ones.

The first effect is the preferential oscillation-induced excitation of the active state. It is based on the state-dependent resonance (strong entrainment in the active state, but not in the background state) in conjunction with supra-linear behavior of the system. The latter could be related either to spike synchronization (like in Schmidt et al. (2018)) or to neural gain function concavity in the subthreshold regime (like in our model). In Schmidt et al. (2018), this effect manifested itself as the ability of high gamma-band oscillations to stabilize an initially metastable active state without affecting the background state. Our results rely on this effect in a more subtle way: the active state in our model is initially stable, but the active module receives additional excitation due to resonant beta-band entrainment, further suppressing the competing background module (which does not get entrained and thus does not receive such excitation).

The second effect is the inhibitory influence of lower-frequency oscillations on the active state, which is related to highly non-linear dynamics that involve pushing the system towards the background state by oscillations. In (Dipoppa, Gutkin, 2013; Schmidt et al., 2018), this effect manifested itself as an oscillation-induced transition to the background state (i.e. erasure of WM content). In both studies, the effect was most prominent in the frequency band just below the resonance of the active state: alpha-band in (Dipoppa, Gutkin, 2013) and beta / low gamma in (Schmidt et al., 2018). In our model, the inhibitory effect of oscillations on the active state is much weaker. Thus, alpha-band oscillations (given beta-band resonance of the active state) do not cause WM erasure by themselves, but play a role in promoting distractor loading: their inhibitory effect on the active state counteracts their excitatory effect discussed above, which allows the distractor-related activity to begin. The WM erasure effect is observed in our model only at very low frequencies (below 1 Hz).

As discussed above, the inhibitory influence of near-resonant oscillations on the active state played the major role in the previous studies, while in our model the excitatory influence dominated. This could be explained by the presence of slow NMDA currents in our model, which makes the active state more robust to brief periods of inhibition (cf. (Tegner et al., 2002)). Another possible explanation is that entrained oscillations in our model are closer to sinusoidal, while in the previous studies they are more pulse-like (due to the rectified input and the supra-threshold regime prone to spike-to-spike synchronization). The inhibitory effect of oscillations seems to depend on the duration of their downward phase, so this effect is presumably stronger when the duty cycle of oscillations is small.

We also note that low frequencies promote a transition from the background to the active state both in our model (alpha-band – loading of weak distractors) and in Schmidt et al. (2018) (delta-band – the “recall” in a single population and the forced random transitions in a competitive noisy multi-population system). However, the underlying effects are different. In Schmidt et al. (2018), the input oscillations are strongly rectified, so one “pulse” of low-frequency oscillations acts as a stimulus presentation, switching a population to the active state. The effect is observed at low frequencies presumably because the “pulse” is long enough in this case. Oscillations in our model are weaker and sinusoidal, so their switching effect on the distractor-selective background module is related to non-linear properties of the system itself and requires simultaneous presentation of the distractor (the entrained oscillations ride on top of the distractor-related excitation). The effect is observed at low frequencies due to their stronger mean excitatory effect on the background module and stronger inhibitory effect on the competing active module. Note also that high-amplitude oscillations in our could be enough to switch one of the modules to the active state without stimulus presentation (given asymmetry in the initial conditions). This effect parallels spontaneous switching between populations induced by delta-band input in the presence of noise reported by Schmidt et al. (2018).

Finally, we should also note that the mechanisms of oscillatory control based on the gain function non-linearity and on the spike synchronization are not mutually exclusive. In the spiking network version of our model, the entrained oscillations that produced the stabilizing and destabilizing effects were much less sinusoidal than in the low-dimensional version. Although we did not analyze this explicitly, we suppose that spike-to-spike synchronization plays a certain role in mediating the effects of oscillations on WM stability in the spiking network version of our model. We suggest that relations and interactions between the aforementioned mechanisms should be explored in more detail in future research.

### General perspectives

#### Separated WM “storage” and “controllers”

We suggest that it could be functionally profitable for the brain to have separate “representational” circuits for retention / processing of information and “controller” circuits that modulate the behavior of the representations. The profit of having a separated controller is especially clear for PFC circuits, given its ability for fast and flexible formation of representations in a task-dependent manner (Parthasarathy et al., 2017; Bouchacourt, Buschman, 2019).

A plausible anatomical substrate for the “representational” circuits is the Layer 2/3 of the neocortex. Circuits of the Layer 2/3 demonstrate sparse firing patterns (Barth, Poulet, 2012), i.e. their neurons likely represent certain features in a highly selective manner. It is also known that the Layer 2/3 networks serve as a basis for WM retention (Goldman-Rakic, 1995). Recently, it was demonstrated that spiking activity in the Layer 2/3 of PFC contains more information about WM content compared to the deeper cortical layers (Bastos et al., 2018).

There are several subcortical structures that may potentially act as a “controller” of retention stability by providing modulatory beta-band signals to the PFC circuits. The first candidate is the basal ganglia (BG), which perform gating function, controlling whether certain information should be placed into WM or ignored (McNab, Klingberg, 2008; Nee, Brown, 2013). It was shown that the temporal profiles of beta activity (with a drop during stimulus presentation and a recovery at the retention period) are similar in the PFC and in the subthalamic nucleus (STN, part of the BG). Importantly, the beta-band activity in both structures was higher during and after distractor presentations compared to presentations of to-be-memorized stimuli (Zavala et al., 2017). Furthermore, beta-band coherence between the STN and the PFC was increased in the case of distractors, suggesting active interaction between these structures (Zavala et al., 2017). Interestingly, the ability to ignore distractors in a WM task is better in patients with Parkinsonian disease (Cools et al., 2010), which is usually accompanied by pathologically increased beta activity in the BG and the cortex (Brittain, Brown, 2014; Pavlides et al., 2015). Another subcortical structure functionally related to the BG and potentially involved in the beta-band control of WM retention is the mediodorsal thalamus (MD). It was shown that beta-band synchrony between the MD and the PFC (with the MD phase-leading the PFC) was increased during WM retention interval and it predicted the accuracy during the task learning, while inhibition of the MD disrupted the synchrony and decreased the accuracy (Parnaudeau et al., 2013).

The subcortical systems that provide alpha-band signals for WM control presumably involve networks of thalamocortical neurons in associative and relay thalamic nuclei, as well as inhibitory circuits of the reticular thalamic nucleus. These structures are known to generate alpha activity in the passive rest condition and participate in shaping top-down attention (Crunelli et al., 2018; Fiebelkorn et al., 2019; Halassa et al., 2014).

It is also likely that the circuits providing oscillatory control of WM are partially located in the deep layers of the neocortex. It is known that the deep cortical layers usually demonstrate LFP oscillations at predominantly low frequencies (Wang, 2010). Among other findings, it was demonstrated that the deep layers during a WM task act as a source of beta activity, which entrains circuits in the superficial layers and presumably controls WM retention (Bastos et al., 2018). Deep cortical layers are tightly interconnected with various subcortical structures, forming thalamocortical and cortico-basal ganglia-thalamocortical loops (Sherman, Guillery, 2018; Shepherd, 2013). Accordingly, deep cortical layers participate in generating beta oscillations in concert with the basal ganglia (Kondabolu et al., 2016; Talakoub et al., 2016; Zavala et al., 2017) and alpha oscillations in concert with the thalamus (Crunelli et al. 2018; Fiebelkorn et al., 2019).

We should note that we do not imply that the mechanisms generating modulatory oscillations are fully non-specific and completely independent of the “representational” circuits’ activity. Instead, we assume that the distribution of various oscillations across the cortex provides a scaffold for the content-specific activity. This scaffold outlines the current task requirements, but it is itself less selective compared to the rate code that represents the WM content. This is in line with lower selectivity of deep cortical layers compared to the superficial ones (Bastos et al., 2018), as well as with lower representational power of subcortical structures compared to the cortex. On the other hand, this oscillatory scaffold is likely refined by feedback projections from the superficial to the deep cortical layers and from the deep cortical layers to subcortical structures. It is clear that WM content retention and its oscillatory modulation are the parts of a unified process, and its self-consistent modeling is an important direction of future research.

#### Oscillatory modulation of encoding capability

We could abstract out from the specific task we modeled in our study (WM with distractors) and consider our results from a more general perspective. We could think of “WM content” as of generic encoded information that is currently being retained or processed by a neural circuit. In this context, one could pose a question about potential ways in which non-specific oscillations (agnostic to the content of the code) could affect the ability of a circuit to retain the encoded information.

In our model, the “code” consists of two binary values: the first value encodes the state of the stimulus-selective module (background or active), and the second value encodes the state of the distractor-selective module. We demonstrated that beta oscillations stabilize the existing code, making it more robust to changes (in agreement with the “beta status quo” hypothesis (Engel, Fries, 2010)), while oscillations below the beta band make the code more prone to changes. From the perspective of cognitive functions, stabilization is required during operations that rely on retention of certain information, such as WM content, task rules, or direction of sustained attention. Conversely, destabilization may be profitable in more “fluent” states such as active perception or divergent thinking.

We also demonstrated that oscillations of very low frequency could erase WM content and prevent memorization (similarly to alpha-band input in (Dipoppa, Gutkin, 2013) and beta-gamma input in (Schmidt et al., 2018)). From the general perspective, such oscillations put the system into a non-coding regime, in which it cannot represent any information. Such regime is profitable when a population should be excluded from ongoing information processing. From the biological perspective, this functionality is similar to the presumable inhibitory role of alpha oscillations (Klimesch, 2012); it is also known that widespread low-frequency oscillations are typical in the states of deep sleep or anesthesia, in which cortical computations are significantly reduced (Akeju, Brown, 2017; Flores et al., 2017).

In summary, our model predicts three distinct functional roles of oscillations in multistable neural systems with population rate coding. First, beta oscillations stabilize the existing code, i.e. make the existing distribution of neural activities harder to change. Second, low-frequency (alpha) oscillations destabilize the code, making it easier to change. Third, oscillations of very low frequency clamp the system into a non-coding state.

## Methods

### General description of the models

In this paper, we used two types of models: (1) spiking networks of leaky integrate-and-fire (LIF) neurons, and (2) low-dimensional population models of Wilson-Cowan type. Individual neurons in the spiking version of the network were coupled by current-based synapses with exponential kernels and no delay. Excitatory currents contained two components with different decay time constants (AMPA and NMDA), while inhibitory currents contained single component (GABAA). Gain functions of the neurons used in the low-dimensional models were numerically precalculated using single-neuron simulations with various input parameters; the precalculated values were then interpolated during network parameter selection and simulations of the low-dimensional models.

We considered both single-module and two-module models. A module always contained an excitatory and an inhibitory population of neurons, with sparse connections within and between the populations. Each population received a noisy background input (explicitly in the spiking models, and implicitly in the low-dimensional models – via an additional term in the equations that describe the dynamics of the AMPA current variance). External oscillations were added to models explicitly as a zero-mean sinusoidal signal affecting the variable that describes the mean AMPA current.

We selected parameters in such way that a module without oscillations demonstrated bistability with realistic firing rates and CV values, as well as beta-band resonance in the active, but not in the background state. During parameter selection, we largely relied on the phase-plane visualization. Then we added input oscillations and simulated low-dimensional models, demonstrating the effects we were interested in. Finally, we confirmed the results by simulating the corresponding spiking models.

#### Spiking models

Each neuron in our spiking models was described by the following equations:

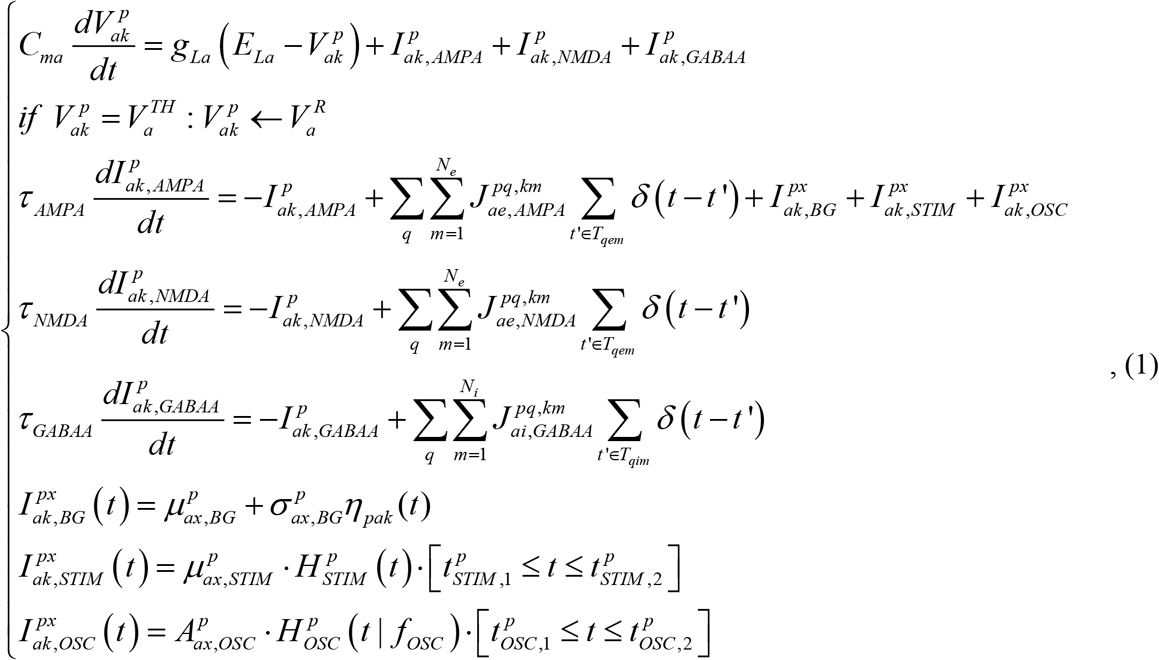

Where *p*, *q* – module indices, *a*, *b* – population types (*a*,*b* = *e* for excitatory and *a*,*b* = *i* for inhibitory populations), *k*, *m* – indices of neurons within populations (the *p*, *a*, *k* indices characterize the neuron, for which the above system of equations is written, while the *q*,*b*, *m* indices characterize the inputs of this neuron); 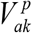 – membrane potential, *E_La_* – resting potential, *C_ma_* – membrane capacitance, *g_La_* – membrane conductance, 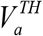 – spike generation threshold, 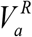 – reset potential; – 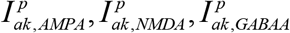 synaptic currents, *τ_AMPA_*,*τ_NMDA_*,*τ_GABAA_* – synaptic time constants, 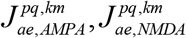 – weights of AMPA and NMDA synaptic connections from the *m* -th neuron of the excitatory population of the module *q* to the *k* -th neuron of the population *a* of the module *p*, 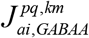 – weight of GABAA synaptic connections from the *m* -th neuron of the inhibitory population of the module *q* to the *k* -th neuron of the population *a* of the module *p*, *T_qem_*,*T_qim_* – time moments of spike generation by the *m* -th neuron of the excitatory / inhibitory population of the module *q*, 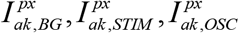 – external current to the *k* -th neuron of the population *a* of the module *p* (BG – background, STIM – representing stimulus presentation, OSC – oscillatory), 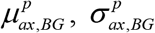 – mean and standard deviation of the background input (same for all neurons of a population), *η_pak_*(*t*) – Gaussian white noise with zero mean and unity variance (independent for each neurons), 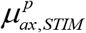 – mean stimulus current, *t*_*STIM*,1_,*t*_*STIM*,2_ time of the stimulus start and end, *H_STIM_*(*t*) – a function defining the stimulus shape; 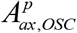, *f*_*OSC*_ – amplitude and frequency of the input oscillations, *H_OSC_*(*t*|*f_osc_*)– a function defining the shape of the oscillations; *t*_*OSC*,1_, *t*_*OSC*,2_ – time of the oscillatory input start and end.

Populations of a type *a* contained *N_a_* neurons, and each of them was connected to *K_ab_* randomly selected neurons from a population of a type *b*. In order to make activities of neurons almost independent, the connectivity was made sparse, i.e. *K_ab_ ≪ N_b_*. Synaptic weights between connected neurons depended only on the populations / modules these neurons belong to:

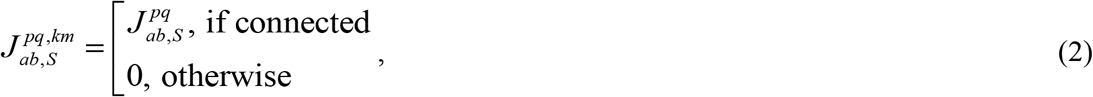

where S=AMPA,NMDA,GABAA.

In the single-module system, we used stimulus of the square shape: 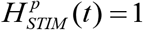. In the two-module system, the stimulus and the distractor had a shape of a square pulse with smoothed fronts, given by the following heuristic formula:

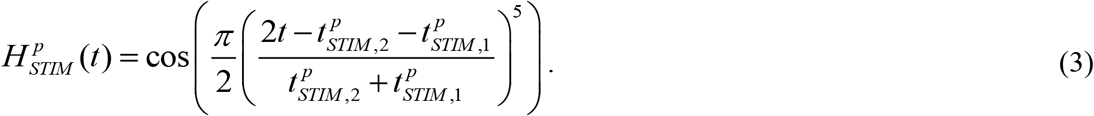

This was motivated by the fact that the phase of oscillations at the end of the distractor presentation could affect the subsequent behavior of the system (the distractor has more chance to be memorized if it terminates at the excitatory phase of oscillations). As we were interested in the effect of oscillations’ frequency (in its relation to the time-averaged firing rates), and not of their phase, we used distractors with smoothed, which allowed us to minimize the phase-dependency effects.

For the oscillatory input, we used a simple sine wave:

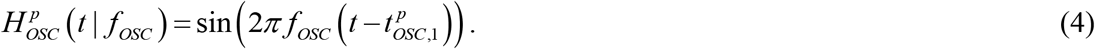

All simulations were performed with the time step Δ*t* = 0.1ms. Instantaneous population firing rate was estimated as the number of spikes generated by population neurons during 1 ms interval, divided by 1 ms and by the number of neurons in the population.

#### Population models

The population models were described by the following system of equations:

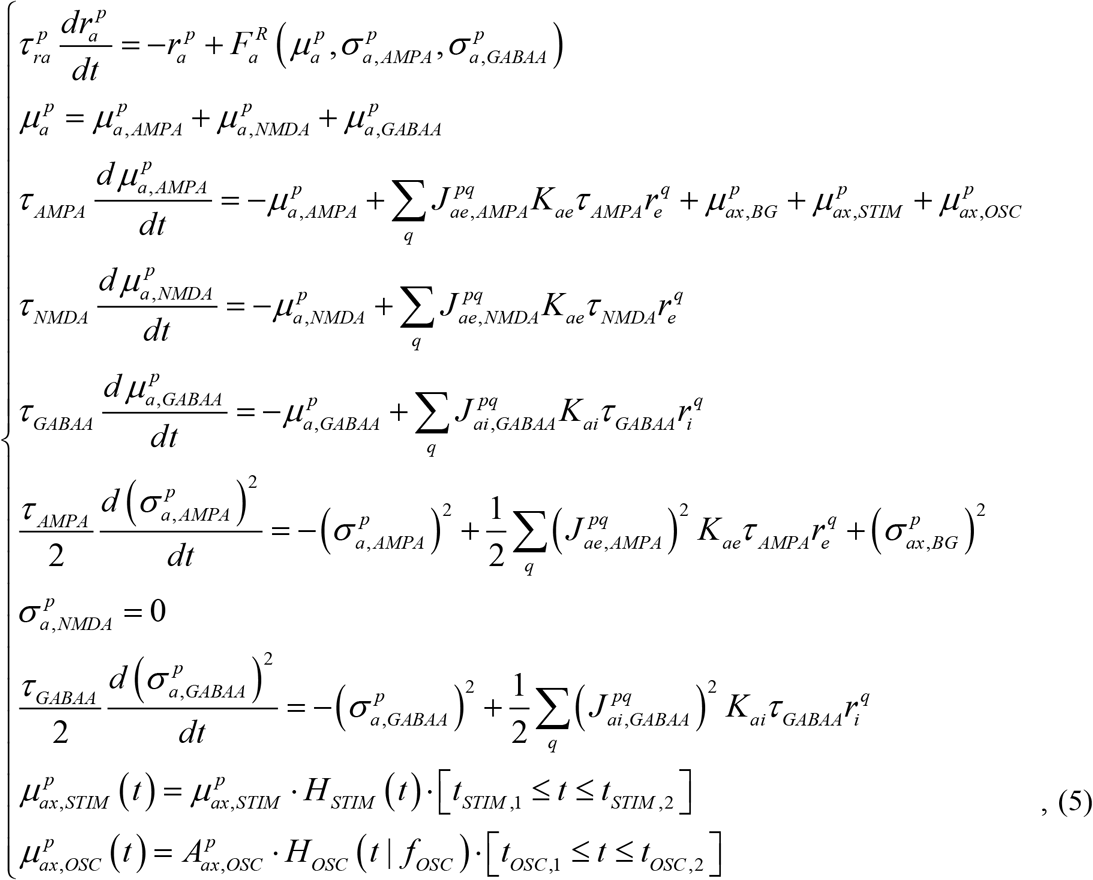

where 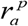 – population frequency (mean number of spikes emitted by a population per unit time, divided by the number of neurons in the population), 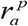 – mean synaptic input current to population neurons, 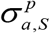 – standard deviation of the synaptic input current over the neurons (S=AMPA,NMDA,GABAA), 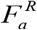 – gain functions which link input mean and standard deviation with output firing rate, *τ_ra_* – time constants of firing rate dynamics; other notations are the same as for the spiking models. We assumed that 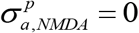, as NMDA receptors are slow, so NMDA-current is almost tonic, with small fluctuations.

Further in the text, we omit the module indices *p*, *q* for most of the parameters, since the modules in the two-module system are identical. For those parameters that are related to the stimulus and the distractor (and, thus, differ between the modules), we use the superscripts “S” and “D”, respectively. For the weights of the inter-module connections, we use the superscript “CROSS”.

We should note that if the low-dimensional system (5) without oscillatory input has steady-states, they quantitatively match quite well the steady-states of the corresponding spiking network (1), given the following conditions: (a) correlation between spike trains of different neurons is low, (b) coefficient of variation of interspike interval (CV) is close to 1 for all neurons, (c) total input firing rate to each neuron is high. However, their dynamical properties (including the response to oscillatory input) could be different, as our low-dimensional model implies fixed time constants τ_ra_, while, in fact, they depend on the state of a spiking network; furthermore, the low-dimensional model does not account for spike-to-spike synchronization. In this work, we do not intend to qualitatively match dynamical behavior of the spiking and the low-dimensional models. Instead, our goal here is to demonstrate that the effects we explore could be observed in both types of models.

#### Precalculation of the gain functions

In order to determine the gain functions 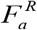 used in (5), we simulated a single LIF neuron receiving a tonic input and two zero-mean noisy inputs with the time constants *τ_AMPA_* and *τ_GABAA_*:

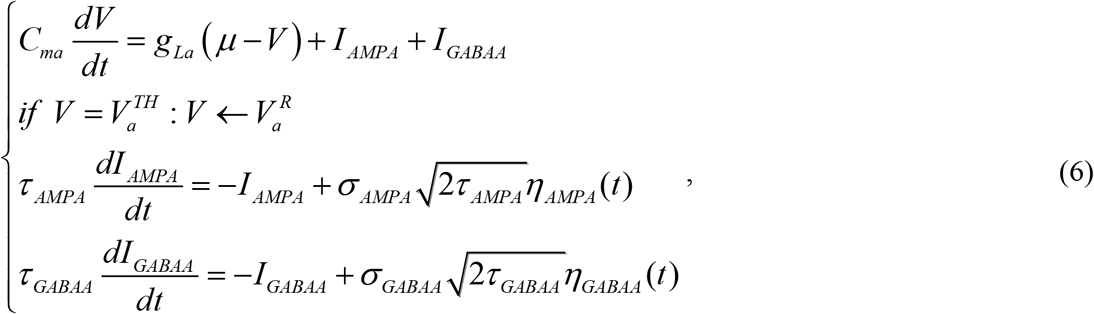

where *μ* corresponds to the depolarization produced by the mean total input current (AMPA + NMDA + GABAA), and *σ_AMPA_,σ_GABAA_* correspond to the total standard deviations of the AMPA- and the GABAA-currents, respectively. We simulated (6) for various combinations (*μ,σ_AMPA_,σ_GABAA_*)located in the nodes of a rectangular grid and we calculated firing rate and CV from the simulated spike trains. Thus, we obtained four 3-D matrices: 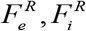 (containing the firing rates) and 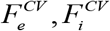 (containing CV values); each position in a matrix corresponded to a certain combination (*μ,σ_AMPA_,σ_GABAA_*).

During parameters selection and simulation of low-dimensional models, we needed to evaluate 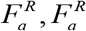 for arbitrary arguments. We did it by sampling appropriate values from the precalculated matrices and applying cubic interpolation to them.

#### Phase-plane analysis

Here we describe the procedure for visualization of the (*r_e_*, *r_i_*) phase plane of the single-module low-dimensional model, with the characteristic *r_e_* - and *r_i_* -curves, such that for each point on the *r_a_* -curve (*a=e* or *i*), derivatives of all the variables except of *r_a_* are equal to zero. Intersections of the *r_e_* – and *r_i_* -curves give the fixed points of the system. In total, the model has 12 dynamical variables: (*r_a_,μ_a,S_,μ_a,GABBA_,(σ_a,S_)*^2^), where *a* = *e*,*i*, *S* = *AMPA*, *NMDA*,*GABAA* for the *μ* -variables, and *S* = *AMPA*,*GABAA* for the σ^2^ -variables. We looked at the manifold in the phase space parametrized by (*r_e_*, *r_i_*)–and defined by the condition that the derivatives of all other variables are zero. This condition could be expressed as follows:

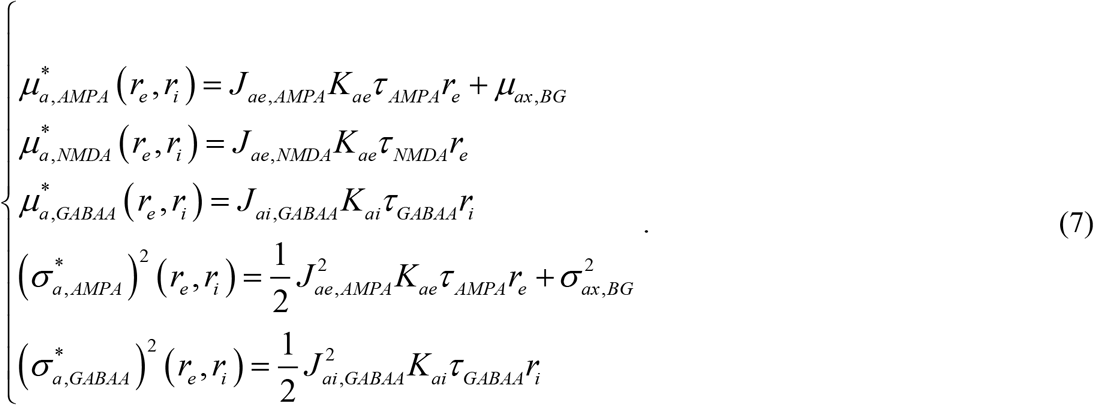

Total mean currents for the points at the manifold could be expressed as follows:

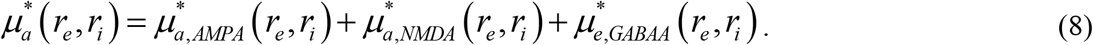

The characteristic *r_e_* - and *r_i_* -curves are defined by the following two conditions, respectively:

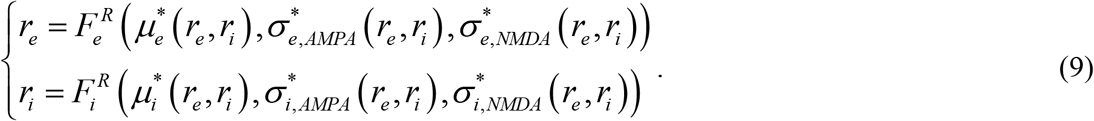

We probed various (*r_e_*, *r_i_*)–combinations from a rectangular grid. For each combination, we applied formulas (7) and (8), and plugged the results into (9). We visualized the characteristic curves as the contour lines, at which the calculated values of 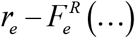(…) and 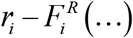(…) changed the sign. Intersections of these curves gave us the fixed points of (5). At the fixed points, we calculated CV values, using a similar interpolation procedure as for the firing rates, but now using the precalculated matrices 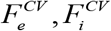. Stability of the fixed points were determined by analyzing the eigenvalues of the Jacobian matrix of (5) calculated at these points.

#### Parameter selection

Out parameter selection procedure could be separated into several steps. First, we selected the parameters of LIF neurons (*C_ma_*,*g_La_*,*E_La_*, 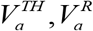) and the synaptic time constants (*τ_AMPA_*,*τ_NMDA_*,*τ_GABAA_*). We chose typical values for these parameters (Table 1) and did not change them for the rest of the process. The excitatory and the inhibitory neurons differed only by the membrane capacitance *C_ma_*; the resulting membrane time constants were: *τ_me_* = *C_me_/g_Le_* = 20 ms and *τ_mi_* = *C_mi_/_gLi_* = 10 ms, which corresponds to pyramidal neurons and fast-spiking interneurons,respectively.

**Table 1.**
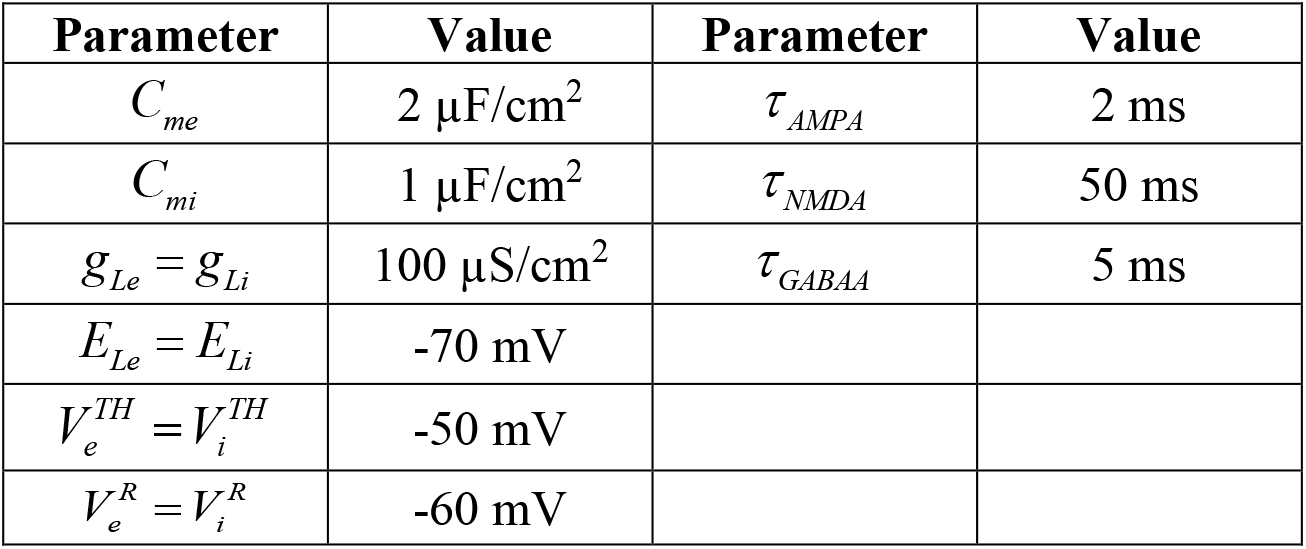
Single-neuron parameters

| Parameter | Value | Parameter | Value |
| --- | --- | --- | --- |
| $C_{me}$ | 2 μF/cm<sup>2</sup> | $\tau_{AMPA}$ | 2 ms |
| $C_{mi}$ | 1 μF/cm<sup>2</sup> | $\tau_{NMDA}$ | 50 ms |
| $g_{Le} = g_{Li}$ | 100 μS/cm<sup>2</sup> | $\tau_{GABAA}$ | 5 ms |
| $E_{Le} = E_{Li}$ | -70 mV | | |
| $V_e^{TH} = V_i^{TH}$ | -50 mV | | |
| $V_e^R = V_i^R$ | -60 mV | | |

Using the selected parameters, we calculated the matrices 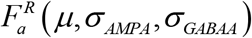 and 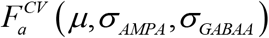. This step was time-consuming, as it involved extensive simulations; however, it was done only once, since all the parameters it requires were selected a priori.

Next, we selected parameters that define steady-state behavior of the single-module system without input oscillations: *K_ab_*,*J_ab,S_*, *μ_ax,BG_*,*σ_ax,BG_*. We chose *K_ee_* = *K_ie_* = 200 and *Kei* = *Kii* = 50. The ratio of inhibitory to excitatory inputs equal to 1:4 is typical; however, the numbers of inputs are about an order of magnitude lower that observed in real cortical neurons. We used them to achieve high sparseness of (*K/N* = 10… 50) without the need to simulate very large networks. We also introduced new parameters *J_ae,TOTAL_* and *k_NMDA_* such that:

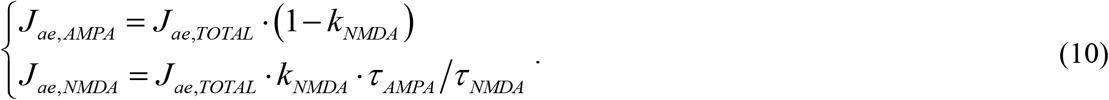

This approach is convenient because *k_NMDA_* only affects the dynamic behavior of the system, but does not affect its fixed points. Our process of *J_ae,TOTAL_*,*J_ai,GABAA_*,*μ_ax,BG_*,*σ_ax,BG_* selection was guided by the phase-plane visualization described in the previous section. We aimed to meet the following requirements: (1) amplitudes of postsynaptic potentials corresponding to selected synaptic weights are in a reasonable range, (2) the system has three fixed points, two of which potentially are steady states (the stability check is described below), (3) both potential steady states have biologically plausible firing rates and CV about 1.

At the next step, we chose the values of *τ_re_*, *τ_ri_*. To do this, we determined the properties of the total input (mean, variance of the AMPA-component, variance of the GABAA-component) that the excitatory and the inhibitory populations receive in the active state (i.e. in the upper of the two steady states that the system has, given the previously selected parameters). For each type of neurons (excitatory and inhibitory), we simulated an uncoupled spiking network that consisted of 1000 LIF neurons receiving an external tonic input equivalent to the total input that the corresponding population received in the active state. We delivered a small current step to the network and averaged the firing rate responses to this step over 1000 trials. Then we fitted an exponential curve to the averaged response for each type of neurons, and the time constants of these exponents were chosen as the values of *τ_re_*,*τ_ri_*.

After that, we checked that two of the fixed points are stable by analyzing the Jacobian matrix of (5) evaluated at each of the fixed points (given zero stimulus-related and oscillatory inputs). Then we selected some intermediate value for *A_ex,OSC_*, and set *A_ix,OSC_* = 0. We obtained the frequency responses of the resulting low-dimensional system by simulating it for various values of *f_osc_*, starting from the background or from the active state. We checked whether the system had a prominent beta-band resonance in the active state and weak resonant properties in the background state. The latter condition was usually met automatically, as the slope of the gain functions (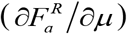) is higher in the active state than in the background state (if CV in both states is close to 1 or higher), so, effectively, interaction between populations is stronger in the active state, which makes the system more resonant. If either the stability condition or the beta-band resonance condition was not satisfied, our first option was to change *k_NMDA_* and to repeat the stability and the frequency response analyses. In general, decrease of *k_NMDA_* shifted the resonant frequency towards higher values and made the resonance stronger; however, too small *k_NMDA_* lead to unstable dynamics with self-generated oscillations (which we wanted to avoid). If changing *k_NMDA_* was not enough to obtain the required properties, we changed *J_ae,TOTAL_*, *J_ai,GABAA_*, *μ_ax,BG_*, guided by the following intuitions: (a) quality of the resonance increases as the system approaches Hopf bifurcation, (b) both quality and frequency of the resonance increase with increasing steady-state firing rate, *J_ee,TOTAL_*, *J_ie,TOTAL_*, and |*J_ei,GABAA_*|, (c) both quality and frequency of the resonance decrease with increasing |*J_ii,GABAA_*|. After changing *J_ae,TOTAL_*, *J_ai,GABAA_*, *μ_ax,BG_*, we recalculated *τ_re_*,*τ_ri_* and repeated the stability and the frequency response analyses.

Next, we repeated the frequency response analysis for various *A_ex,OSC_*. Our goal was to find *A_ex,OSC_*, for which a beta-band input would selectively produce a prominent shift of the time-averaged firing rate in the active state, but not in the background state. To achieve a strong shift, *A_ex,OSC_* should be high enough so non-linearity of the gain functions plays a significant role. On the other hand, we tried to stay close to physiological regime of oscillations and avoid excessively synchronized and rectified all-or-none dynamics. If we failed to achieve the required behavior, we changed *k_NMDA_* or *J_ae,TOTAL_*,*J_ai,GABAA_*, *μ_ax,BG_*, repeated the required steps, and returned to this analysis. This was the final step of the low-dimensional single-module system configuration.

After configuring the low-dimensional model, we used the selected parameters for simulating a spiking network of *N* =10000 neurons. First, we noted that the firing rates corresponding to the active state were higher in the spiking network than in the low-dimensional model (probably due to finite-size effects or due to fluctuations of NMDA current, not accounted for in the low-dimensional model); thus, we selected slightly smaller *μ_ex,BG_* value for the spiking model, leaving other parameters unchanged. Second, we found that the spiking model responded to a strong oscillatory input by excessive synchronization; this also led to a shift of the resonant peak from the beta band towards lower frequencies. To treat this issue, we selected the ratio *A_ix,OSC_* : *A_ex,OSC_* = 0.2 (instead of *A_ix,OSC_* = 0 in the low-dimensional model) and compensated it with an increase of *A_ex,OSC_*.

The two-module system consisted of two identical modules with symmetrical mutual inhibition implemented via connections from the excitatory population of each module to the inhibitory population of another module. To compensate for this additional inhibition, we slightly increased *μ_ex,BG_* (compared to the single-module system) for each of the two modules. The spiking version of the model contained *N* = 5000 neurons in each module.

Amplitude of the stimulus was chosen to be enough to switch the receiving module to the active state, whether the other module is in the background or in the active state. Amplitude of the strong distractor was equal to the stimulus amplitude. Amplitude of the weak distractor was decreased in such way that the distractor presentation did not change the state of the system.

Parameters of the single-module system are presented in Table 2. Those parameters of the two-module system that differ from the single-module system are presented in Table 3.

**Table 2.**
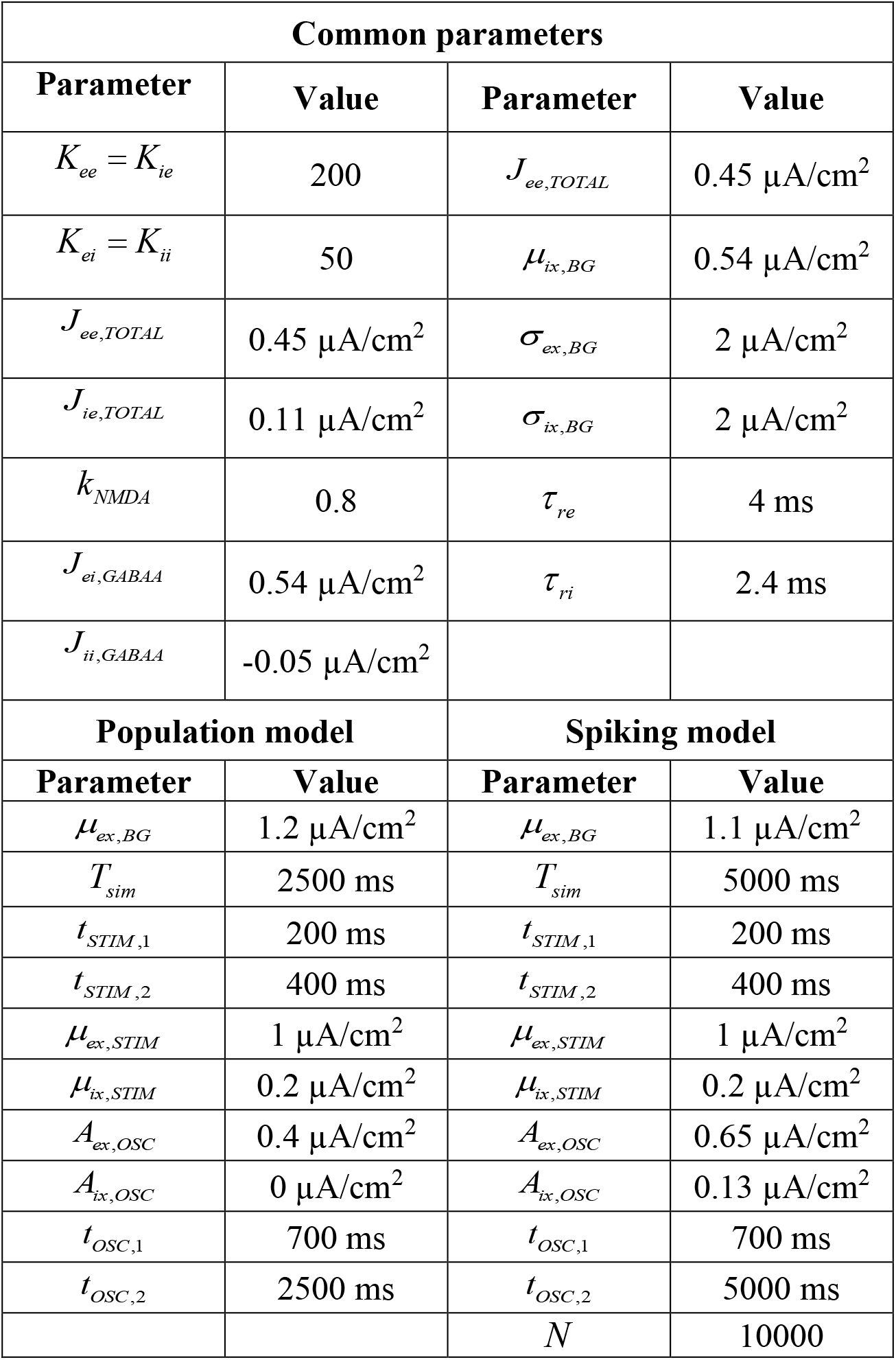
Parameters of the single-module system

| Common parameters |  |  |  |
| --- | --- | --- | --- |
| Parameter | Value | Parameter | Value |
| $K_{ee} = K_{ie}$ | 200 | $J_{ee,TOTAL}$ | $0.45 \mu\text{A}/\text{cm}^2$ |
| $K_{ei} = K_{ii}$ | 50 | $\mu_{ix,BG}$ | $0.54 \mu\text{A}/\text{cm}^2$ |
| $J_{ee,TOTAL}$ | $0.45 \mu\text{A}/\text{cm}^2$ | $\sigma_{ex,BG}$ | $2 \mu\text{A}/\text{cm}^2$ |
| $J_{ie,TOTAL}$ | $0.11 \mu\text{A}/\text{cm}^2$ | $\sigma_{ix,BG}$ | $2 \mu\text{A}/\text{cm}^2$ |
| $k_{NMDA}$ | 0.8 | $\tau_{re}$ | 4 ms |
| $J_{ei,GABAA}$ | $0.54 \mu\text{A}/\text{cm}^2$ | $\tau_{ri}$ | 2.4 ms |
| $J_{ii,GABAA}$ | $-0.05 \mu\text{A}/\text{cm}^2$ | | |
| Population model |  | Spiking model |  |
| Parameter | Value | Parameter | Value |
| $\mu_{ex,BG}$ | $1.2 \mu\text{A}/\text{cm}^2$ | $\mu_{ex,BG}$ | $1.1 \mu\text{A}/\text{cm}^2$ |
| $T_{sim}$ | 2500 ms | $T_{sim}$ | 5000 ms |
| $t_{STIM,1}$ | 200 ms | $t_{STIM,1}$ | 200 ms |
| $t_{STIM,2}$ | 400 ms | $t_{STIM,2}$ | 400 ms |
| $\mu_{ex,STIM}$ | $1 \mu\text{A}/\text{cm}^2$ | $\mu_{ex,STIM}$ | $1 \mu\text{A}/\text{cm}^2$ |
| $\mu_{ix,STIM}$ | $0.2 \mu\text{A}/\text{cm}^2$ | $\mu_{ix,STIM}$ | $0.2 \mu\text{A}/\text{cm}^2$ |
| $A_{ex,OSC}$ | $0.4 \mu\text{A}/\text{cm}^2$ | $A_{ex,OSC}$ | $0.65 \mu\text{A}/\text{cm}^2$ |
| $A_{ix,OSC}$ | $0 \mu\text{A}/\text{cm}^2$ | $A_{ix,OSC}$ | $0.13 \mu\text{A}/\text{cm}^2$ |
| $t_{OSC,1}$ | 700 ms | $t_{OSC,1}$ | 700 ms |
| $t_{OSC,2}$ | 2500 ms | $t_{OSC,2}$ | 5000 ms |
| | | $N$ | 10000 |

**Table 3.**
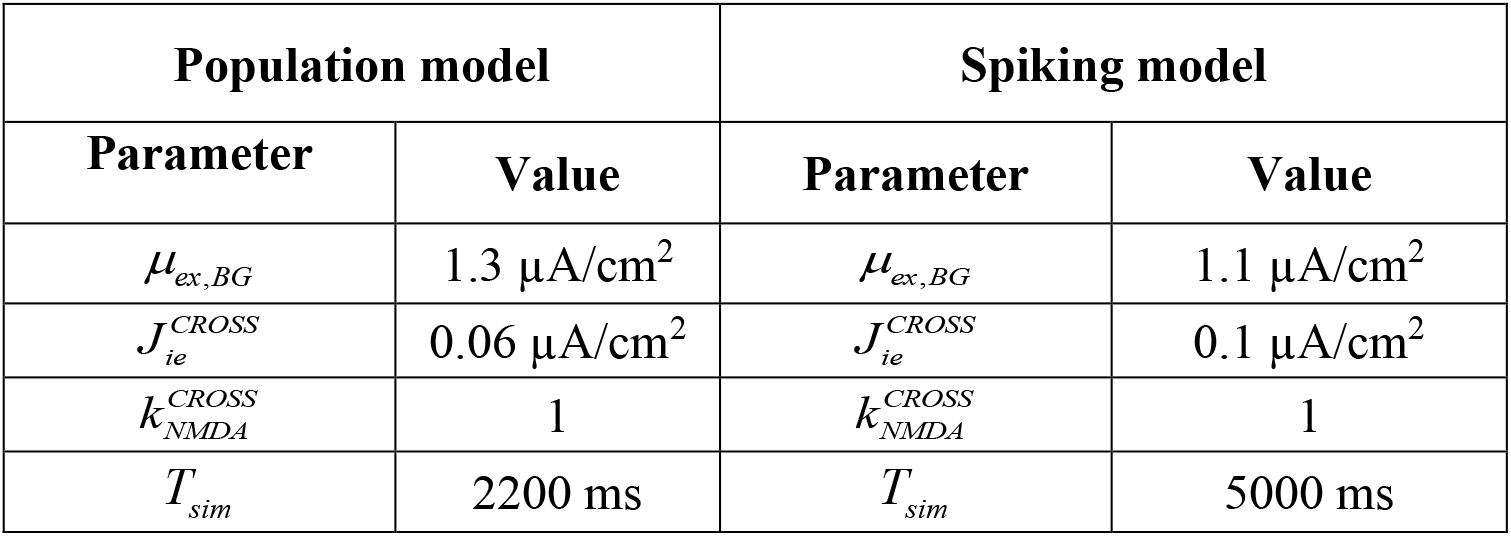
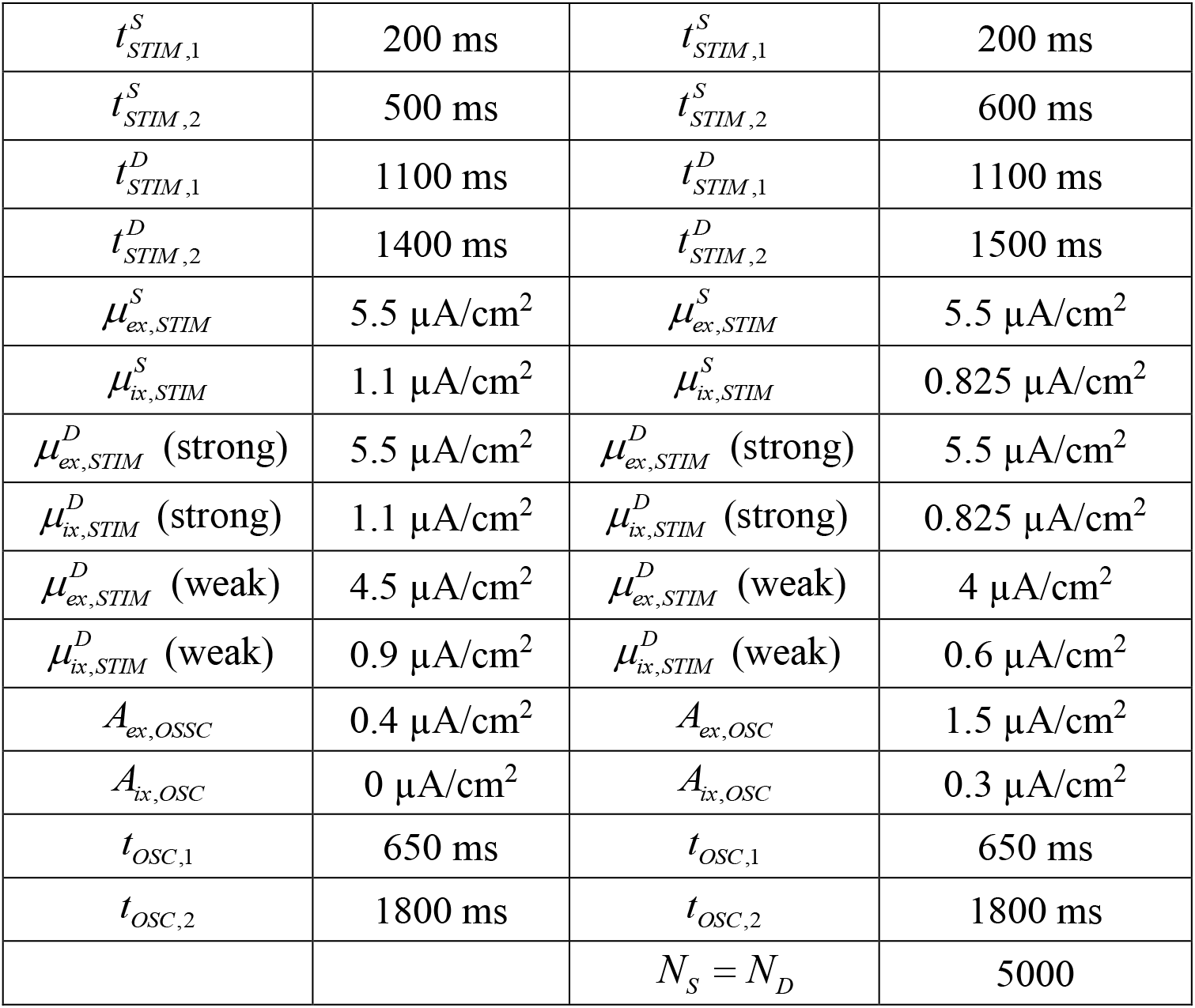
Parameters of the two-module system

| Population model |  | Spiking model |  |
| --- | --- | --- | --- |
| Parameter | Value | Parameter | Value |
| $\mu_{ex,BG}$ | $1.3 \mu\text{A}/\text{cm}^2$ | $\mu_{ex,BG}$ | $1.1 \mu\text{A}/\text{cm}^2$ |
| $J_{ie}^{CROSS}$ | $0.06 \mu\text{A}/\text{cm}^2$ | $J_{ie}^{CROSS}$ | $0.1 \mu\text{A}/\text{cm}^2$ |
| $k_{NMDA}^{CROSS}$ | 1 | $k_{NMDA}^{CROSS}$ | 1 |
| $T_{sim}$ | 2200 ms | $T_{sim}$ | 5000 ms |
| $t_{STIM,1}^S$ | 200 ms | $t_{STIM,1}^S$ | 200 ms |
| $t_{STIM,2}^S$ | 500 ms | $t_{STIM,2}^S$ | 600 ms |
| $t_{STIM,1}^D$ | 1100 ms | $t_{STIM,1}^D$ | 1100 ms |
| $t_{STIM,2}^D$ | 1400 ms | $t_{STIM,2}^D$ | 1500 ms |
| $\mu_{ex,STIM}^S$ | 5.5 $\mu\text{A}/\text{cm}^2$ | $\mu_{ex,STIM}^S$ | 5.5 $\mu\text{A}/\text{cm}^2$ |
| $\mu_{ix,STIM}^S$ | 1.1 $\mu\text{A}/\text{cm}^2$ | $\mu_{ix,STIM}^S$ | 0.825 $\mu\text{A}/\text{cm}^2$ |
| $\mu_{ex,STIM}^D$ (strong) | 5.5 $\mu\text{A}/\text{cm}^2$ | $\mu_{ex,STIM}^D$ (strong) | 5.5 $\mu\text{A}/\text{cm}^2$ |
| $\mu_{ix,STIM}^D$ (strong) | 1.1 $\mu\text{A}/\text{cm}^2$ | $\mu_{ix,STIM}^D$ (strong) | 0.825 $\mu\text{A}/\text{cm}^2$ |
| $\mu_{ex,STIM}^D$ (weak) | 4.5 $\mu\text{A}/\text{cm}^2$ | $\mu_{ex,STIM}^D$ (weak) | 4 $\mu\text{A}/\text{cm}^2$ |
| $\mu_{ix,STIM}^D$ (weak) | 0.9 $\mu\text{A}/\text{cm}^2$ | $\mu_{ix,STIM}^D$ (weak) | 0.6 $\mu\text{A}/\text{cm}^2$ |
| $A_{ex,OSSC}$ | 0.4 $\mu\text{A}/\text{cm}^2$ | $A_{ex,OSC}$ | 1.5 $\mu\text{A}/\text{cm}^2$ |
| $A_{ix,OSC}$ | 0 $\mu\text{A}/\text{cm}^2$ | $A_{ix,OSC}$ | 0.3 $\mu\text{A}/\text{cm}^2$ |
| $t_{OSC,1}$ | 650 ms | $t_{OSC,1}$ | 650 ms |
| $t_{OSC,2}$ | 1800 ms | $t_{OSC,2}$ | 1800 ms |
| | | $N_S = N_D$ | 5000 |

## Acknowledgements

This work was supported by HSE Basic Research Program and the Russian Academic Excellence Project “5100”. BSG acknowledge partial support from CNRS, INSERM, ANR-17-EURE-0017 and ANR-10-IDEX-0001-02.

